# Aldh3a1-mediated detoxification of reactive aldehydes contributes to distinct muscle responses to Amyotrophic Lateral Sclerosis progression

**DOI:** 10.1101/2024.12.02.626422

**Authors:** Ang Li, Li Dong, Xuejun Li, Jianxun Yi, Jianjie Ma, Jingsong Zhou

**Affiliations:** Department of Kinesiology, College of Nursing and Health Innovation, University of Texas at Arlington, TX, 76019, USA; Department of Surgery, Division of Surgical Sciences, University of Virginia, Charlottesville, VA, 22903, USA

**Keywords:** Amyotrophic lateral sclerosis, skeletal muscle, extraocular muscle, reactive aldehydes, aldehyde dehydrogenase, sciatic nerve transection

## Abstract

Different muscles exhibit varied susceptibility to degeneration in Amyotrophic Lateral Sclerosis (ALS), a fatal neuromuscular disorder. Extraocular muscles (EOMs) are particularly resistant to ALS progression, and exploring the underlying molecular nature may offer significant therapeutic value. Reactive aldehyde 4-hydroxynonenal (HNE) is implicated in ALS pathogenesis, and Aldh3a1 is an inactivation-resistant intracellular aldehyde dehydrogenase that detoxifies 4-HNE to protect eyes against UV-induced oxidative stress. We detected prominently higher levels of *Aldh3a1* in mouse EOMs compared to other muscles under normal physiological conditions. In an ALS mouse model (hSOD1^G93A^) reaching end-stage, *Aldh3a1* expression was maintained high in EOMs, substantially elevated in soleus and diaphragm, but only moderately increased in extensor digitorum longus (EDL) muscle, which endured the most severe pathological remodeling, as demonstrated by unparalleled upregulation of a denervation marker *Ankrd1*. Importantly, sciatic nerve transection in wildtype mice further confirmed induced *Aldh3a1* and *Ankrd1* expression in an inverse manner across muscle types in response to denervation. Mechanistically, whole-muscle RNA-Seq and pharmacological tests indicate that higher basal levels of lipid oxidation in soleus and diaphragm muscles may predispose them to stronger Nrf2 antioxidant responses under pathological stress compared to EDL, leading to more prominent *Aldh3a1* upregulation. Additionally, the identification of the myoblast fusion marker *Mymk* as an EOM signature gene suggests that the spontaneous activation of satellite cells contributes to high levels of *Aldh3a1* in EOMs. Functionally, adeno-associated virus-mediated overexpression of *Aldh3a1* protected myotubes from 4-HNE-induced DNA fragmentation and plasma membrane leakage. It also restored MG53-mediated membrane repair, highlighting its potential for clinical applications.

## Introduction

Amyotrophic Lateral Sclerosis (ALS) is a devastating neuromuscular disorder characterized by progressive motor neuron death and skeletal muscle wasting. Remarkably, extraocular muscles (EOMs) exhibit superior preservation of structure, neuromuscular junction (NMJ) integrity and function in ALS patients and rodent models [1–5]. Beside EOMs, animal model studies also show that limb and body muscles respond to ALS progression differently, with fast-twitch muscles generally more susceptible to degeneration than in slow-twitch muscles [6–8]. Factors underlying this phenomenon can be multi-faceted and could hold clues for identifying novel therapeutic targets against muscle degeneration in ALS or other neuromuscular disorders.

Reactive aldehydes resulting from oxidative stress are involved in the pathological process of multiple neurodegenerative disorders including ALS [9]. One major route for intracellular production of reactive aldehydes is the peroxidation of polyunsaturated fatty acids (PUFA), which are major component of cellular membranous structures [10, 11]. Well-known examples include 4- hydroxynonenal (4-HNE), malondialdehyde (MDA) and acrolein [11], which can form adducts with protein through Michael addition or Schiff base formation [12–14]. These adducts contribute to protein crosslinking and aggregation [11], leading to broad pathological consequences including disrupted cell signaling, altered gene expression, inhibited enzyme activity, compromised mitochondrial function and disformed cytoskeleton [15]. Reactive aldehydes can also form adducts with DNA, resulting in inter-strand crosslinks, base substitution, mutation and fragmentation [16–22]. Furthermore, 4-HNE is known to form adducts with phosphatidylethanolamine and certain ion channel proteins at the plasma membrane, causing altered ionic homeostasis and cellular osmolarity [23, 24]. The deprivation of local antioxidants like glutathione by 4-HNE can promote further lipid peroxidation at the plasma membrane, creating a vicious cycle that leads to cell death [25, 26]. Elevated levels of lipid peroxidation markers, including 4-HNE adducts, have been detected in cells of central nervous system and body fluids from patients and/or models of amyotrophic lateral sclerosis (ALS), Alzheimer disease (AD), Parkinson disease (PD) and Huntington disease (HD) [9].

To neutralize reactive aldehydes, the human body has deployed a series of detoxification enzymes. Aldehyde dehydrogenases (ALDHs) aimed to oxidize the carbonyl group into corresponding acids, while aldo-keto reductases (AKRs) reduce aldehydes into corresponding alcohols [12]. Aldh2, Aldh1a1 and Aldh3a1 are the three most studied ALDHs [27]. Aldh2 is a mitochondrial ALDH abundantly present in liver, brain, heart and muscle [28]. Aldh1a1 and Aldh3a1 are primarily cytosolic and are extremely abundant in mammalian corneal and lens to protect against ultraviolet radiation (UVR) induced generation of reactive aldehydes and their pathological consequences [20, 29, 30]. Aldh3a1 knockout mice and Aldh1a1/Aldh3a1 double knockout mice develop cataracts by 1 month of age [30]. It is also worth noticing that a kinetics study of these three ALDHs reported that Aldh2 was irreversibly inactivated by 4-HNE and acrolein at above 10 μM. Aldh1a1 was inactivated by acrolein at concentrations higher than 1 mM, while no inactivation of Aldh3a1 was observed by either 4-HNE, acrolein or MDA even at 20 mM [27]. Thus, Aldh3a1 is the most inactivation-resistant isoform of the three and the focus of the current study.

It is unknown whether aldehyde dehydrogenases are involved in varied susceptibility of different muscles to degeneration under ALS. In this study, we examined the expression of *Aldh2*, *Aldh1a1* and *Aldh3a1* in EDL, soleus, diaphragm and EOMs of end-stage hSOD1^G93A^ (G93A) mice, a well-established ALS rodent model, as well as their wildtype (WT) littermates [31]. Only *Aldh3a1* exhibited dramatically higher expression in EOMs compared to other muscles in WT mice. Meanwhile it was prominently upregulated in G93A soleus and diaphragm compared to WT controls. However, the upregulation was less pronounced in G93A EDL muscle, which suffers the most severe NMJ degeneration in these four muscle types in G93A mice [5]. Indeed, the commonly used denervation marker *Ankrd1* [32, 33] was induced most in G93A EDL muscle, suggesting that the expression level of *Aldh3a1* is inversely linked to the severity of muscle pathological remodeling. The distinct expression pattern of *Aldh3a1* gene was also confirmed at the protein level. In EDL and soleus muscles of WT mice with sciatic nerve transection (SNT), *Ankrd1* expression was quickly elevated post operation, especially in EDL muscles and gradually decreased over time. In contrast, the upregulation of *Aldh3a1* occurred with a multi-day delay in soleus, while no significant upregulation of *Aldh3a1* occurred in EDL muscles even after 14 days. Thus, *Ankrd1* and *Aldh3a1* exhibit inverse induction pattern over muscle type and time post denervation.

Whole muscle RNA-seq analysis and pharmacological tests revealed potential mechanisms underlying the differential regulations of *Aldh3a1* expression in various muscles involving Nrf2 signaling activation. Importantly, transduction of EDL and soleus-SC derived myotubes with adeno-associated virus (AAV) vector expressing human *Aldh3a1* markedly decreased 4-HNE-induced DNA fragmentation. Furthermore, MG53 is a muscle specific tripartite motif family protein nucleating the assembly of the repair machinery on injured plasma membrane [34]. We previously reported abnormal MG53 intracellular aggregation and compromised membrane repair in ALS muscles [35]. Here, we demonstrated that enforced expression of *Aldh3a1* in cultured myotubes protected against 4-HNE-induced plasma membrane damage and restored MG53 mediated membrane repair. Thus, the differential expression level of *Aldh3a1* in muscles in response to ALS progression likely reflects a varying protective role of Aldh3a1 against reactive aldehyde cytotoxicity in various muscles. Organ-targeted delivery of Aldh3a1 through gene therapy may add a new therapeutic avenue for ALS or diseases associated with elevated oxidative stress.

## Results

### Differential expression of *Aldh3a1* across muscles and ALS disease states

The transcription levels of three aldehyde dehydrogenases (*Aldh1a1*, *Aldh2* and *Aldh3a1*) in muscles from different anatomic origins, including hindlimb (EDL and soleus), diaphragm and EOMs were evaluated by qRT-PCR. These muscles were collected from WT mice and their end-stage G93A littermates (4 pairs of male mice and 4 pairs of female mice at 4-5 months of age). As shown in **Figure 1A** left panels and **Figure 1-figure supplement 1A**, *Aldh3a1* was expressed prominently higher in EOMs than other muscles in WT mice (59-fold that in EDL, 22-fold that in soleus and 91 folds that in diaphragm). Notable induction of *Aldh3a1* was detected for soleus (50-fold) and diaphragm (79-fold) from G93A mice compared to WT controls, while the induction was far less pronounced for EDL (7-fold). In G93A EOMs, *Aldh3a1* expression was maintained at comparable levels to WT EOMs. This pathological induction pattern is interesting as EDL is the most severely affected during ALS progression [4, 5]. Importantly, denervation-associated pathological remodeling marker *Ankrd1* (*CARP)* exhibited stronger induction in G93A EDL (69-fold) than diaphragm (35-fold), soleus (22-fold) and EOMs (3-fold) (**Figure 1A** right panels and **Figure 1-figure supplement 1B**). Data here suggests a potential relationship between *Aldh3a1* expression and muscle resistance to degeneration in ALS. In contrast to *Aldh3a1*, neither *Aldh1a1* nor *Aldh2* exhibited superior expression in EOMs, and none of the G93A muscles exhibited differential expression of *Aldh1a1* compared to WT controls, implying that it is unlikely involved in muscle dependent response to ALS progression (**Figure 1-figure supplementary 1C,D** and **Figure 1-Source Data 1**). Thus, *Aldh3a1* is the focus of this study.

**Figure 1.**
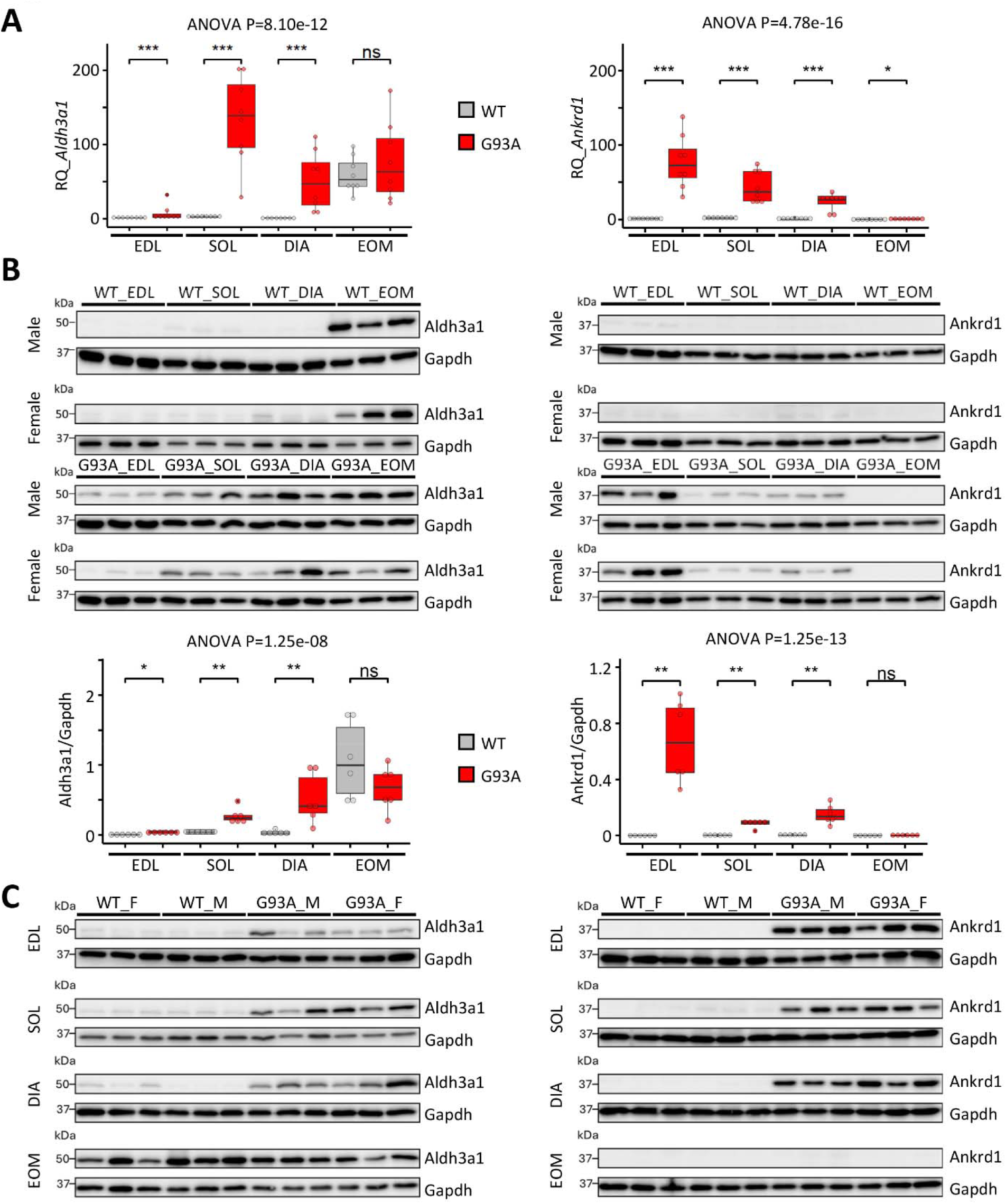
Differential expression of *Aldh3a1* and *Ankrd1* in different muscles from end-stage G93A mice and WT littermates. (**A**) Comparing RNA levels of *Aldh3a1* and *Ankrd1* in Extensor Digitorum Longus (EDL), soleus (SOL), diaphragm (DIA) to those in extraocular muscles (EOMs) by qRT-PCR. N = 8 (4 pairs of male and 4 pairs of female). RQ, relative quantification (WT EDL samples served as the reference group). *** P < 0.001; * P < 0.05; ns, not significant (Wilcoxon rank-sum test). ANOVA P values are also shown. Please also see **Figure 1-figure supplement 1** and **Figure 1-Source Data 1**. (**B**) Western blots of Aldh3a1, Ankrd1 and housekeeping protein Gapdh in different muscles from end-stage G93A mice and WT littermates. The quantification results used WT EDL samples as the reference group. N = 6 (3 pairs of male and 3 pairs of female). * P < 0.05; ** P < 0.01; ns, not significant (Wilcoxon rank-sum test). ANOVA P values are also provided. Please also see **Figure 1-figure supplement 2** and **Figure 1-Source Data 2**. (**C**) Western blots of Aldh3a1, Ankrd1 and housekeeping protein Gapdh in the same muscle from end-stage G93A mice and WT littermates. F: female, M: male.

The unique expression pattern of *Aldh3a1* is also present at the protein level. In WT mice, Aldh3a1 protein was most abundant in EOMs (132-fold that in EDL, 24-fold that in soleus and 29-fold that in diaphragm after normalization to housekeeping protein Gapdh). In line with the qPCR results, the abundance of Aldh3a1 protein was maintained at high levels in EOMs in end-stage G93A mice. Remarkably, significantly elevated levels of Aldh3a1 protein were detected in diaphragm (14-fold), soleus (6-fold) and EDL muscle (5-fold) compared to WT counterparts (**Figure 1B, C, Figure 1-figure supplement 1A, 2** and **Figure 1-Source Data 2**). In comparison, Ankrd1 protein increased most significantly in G93A EDL muscle compared to WT counterparts (594-fold), followed by diaphragm (66-fold), soleus (51-fold) (**Figure 1B, C**, **Figure 1-figure supplement 1B, 2** and **Figure 1-Source Data 2**). Consistently, in the transverse section of EDL muscles from G93A mice, most myofibers were positively stained with anti-Ankrd1 antibodies (**Figure 2A**). Ankrd1 positive myofibers were relatively sparse in G93A soleus and diaphragm muscles, while in G93A EOMs they were extremely scarce. To further validate the association between Ankrd1 and denervation, we performed sciatic nerve transection (SNT) in the right hindlimbs of WT mice 4-5 months of age. The majority of EDL and soleus myofibers became Ankrd1 positive post SNT (**Figure 2B**). Thus, our data implied an inverse relationship between *Aldh3a1* expression pattern and that of the denervation marker *Ankrd1* in G93A muscles.

**Figure 2.**
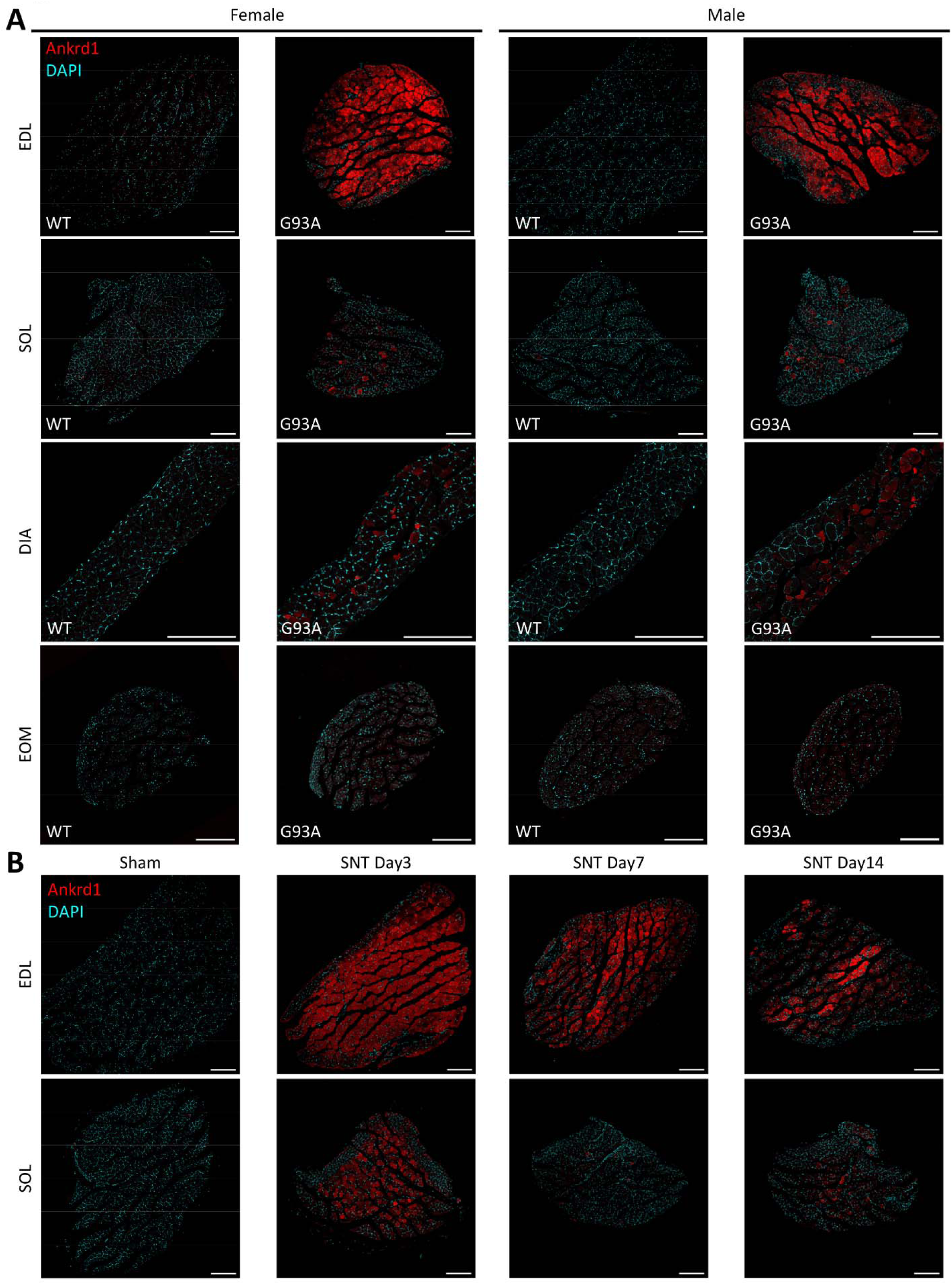
Global view of Ankrd1 immunostaining patterns in transverse sections of different muscles. (**A**) Transverse sections of different muscles from end-stage G93A mice and WT controls stained with anti-Ankrd1 antibodies and counterstained with DAPI. N = 6 (3 pairs of male and 3 pairs of female). Scale bars, 200 μm. (**B**) Transverse sections of EDL and soleus muscles dissected from WT mice (4-5 months of age) at Day 3, Day 7 and Day 14 post sciatic nerve transection and sham operated controls stained with Ankrd1 antibodies and counterstained with DAPI. N = 6 (3 male and 3 female). Scale bars, 200 μm.

We did whole-mount immunostaining to investigate the distribution of Aldh3a1 protein in myofibers (**Figure 3A** and **Figure 3-figure supplement 1A**). In WT mice, EOM exhibited higher levels of cytosolic Aldh3a1 than other muscles, confirming the Western blot and qPCR data in **Figure 1**. In muscles from end-stage G93A mice, soleus and diaphragm show increased Aldh3a1 in the cytosol and myonuclei, while Aldh3a1 positive myofibers were scarce in EDL (**Figure 3A** and **Figure 3-figure supplement 1A**, yellow arrows). Additionally, axon terminals (labelled by Sg2 antibody) were partially or fully absent from AChR positive area (labelled by BTX), confirming partial or full denervation of G93A muscles. These data further support that the induction of *Aldh3a1* expression is less robust in G93A EDL than in soleus and diaphragm.

**Figure 3.**
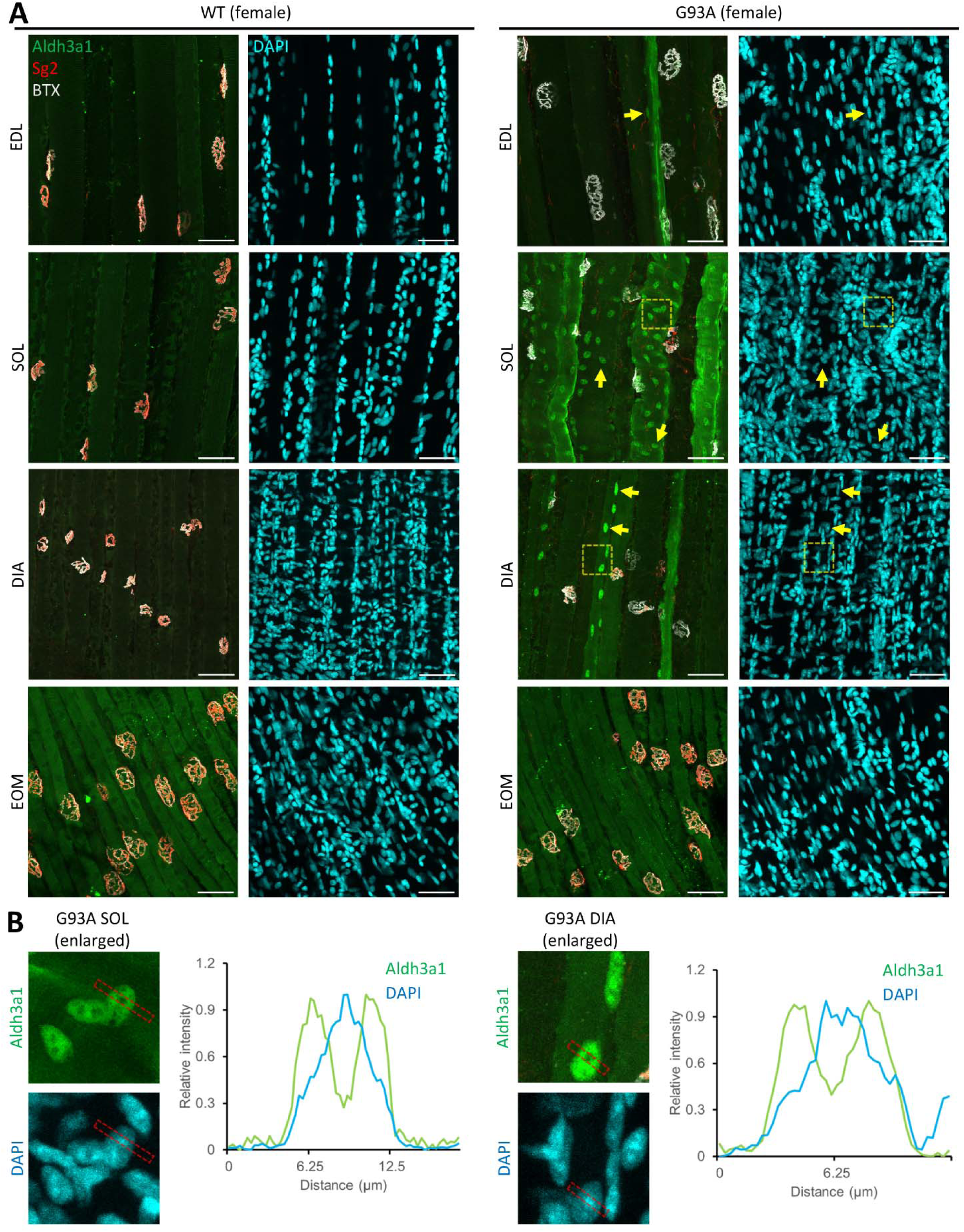
Whole-mount immunostaining of Aldh3a1 in different muscles from end-stage G93A mice and WT littermates. (**A**) Representative compacted z-stack scan images of whole-mount EDL, soleus diaphragm extraocular muscles stained with antibodies against Aldh3a1, Sg2 (labeling axon terminals), Alexa Fluor conjugated α-Bungarotoxin (BTX, labeling AChRs on muscle membrane) and DAPI (labeling nuclei). Yellow arrows highlight nuclei with Aldh3a1 enrichment. Dashed yellow boxes denote regions enlarged in Panel B for kymographic measurement. N = 6 (3 pairs of male and 3 pairs of female). Scale bars, 50 μm. Please also see **Figure 3-figure supplement 1A**. (**B**) Profiling the relative intensity of Aldh3a1 and DAPI fluorescent signals along the strips denoted by dashed red boxes. Relative intensities are calculated as (*F* − *F_min_*)/(*F_max_* − *F_min_*). Also see **Figure 3-figure supplement 1B**.

It is worth noticing that the nuclear Aldh3a1 signal dips in the DAPI dense foci, where genomic DNA is highly compacted (**Figure 3B** and **Figure 3-figure supplement 1B**) [36], implying that nuclear Aldh3a1 preferentially localizes to the euchromatin regions, a pattern commonly seen for histone modifications associated with actively transcribing genes [37]. We speculate that the elevation of Aldh3a1 levels in G93A muscles is a self-defense mechanism protecting cytosolic protein and vulnerable genomic DNA regions against lipid peroxidation stress under ALS.

We also examined whether there is an inverse correlation between aldh3a1 and Ankrd1 distribution on the level of individual myofibers. Most myofibers in G93A EDL were positive for Ankrd1, with few of them also positive for Aldh3a1 (**Figure 4A**, yellow arrows). Ankrd1 positive myofibers were sparser in G93A soleus and diaphragm, whereas myofibers with elevated cytosolic and nuclear Aldh3a1 were more frequently seen (**Figure 4B, C**, yellow arrows). These Aldh3a1 positive myofibers could either be Ankrd1 positive (yellow arrows) or negative (blue arrows) (**Figure 4B, C**). In G93A EOMs, Ankrd1 positive myofibers were extremely scarce, whereas most G93A EOM myofibers exhibited cytosolic distribution of Aldh3a1 (**Figure 4D**), a feature shared by their WT peers (**Figure 4-figure supplement 1**). In sum, although Aldh3a1 and Ankrd1 are not mutually exclusive in individual myofibers, their abundance does exhibit an inverse relationship on the whole-muscle scale across different muscles in end-stage G93A mice.

**Figure 4.**
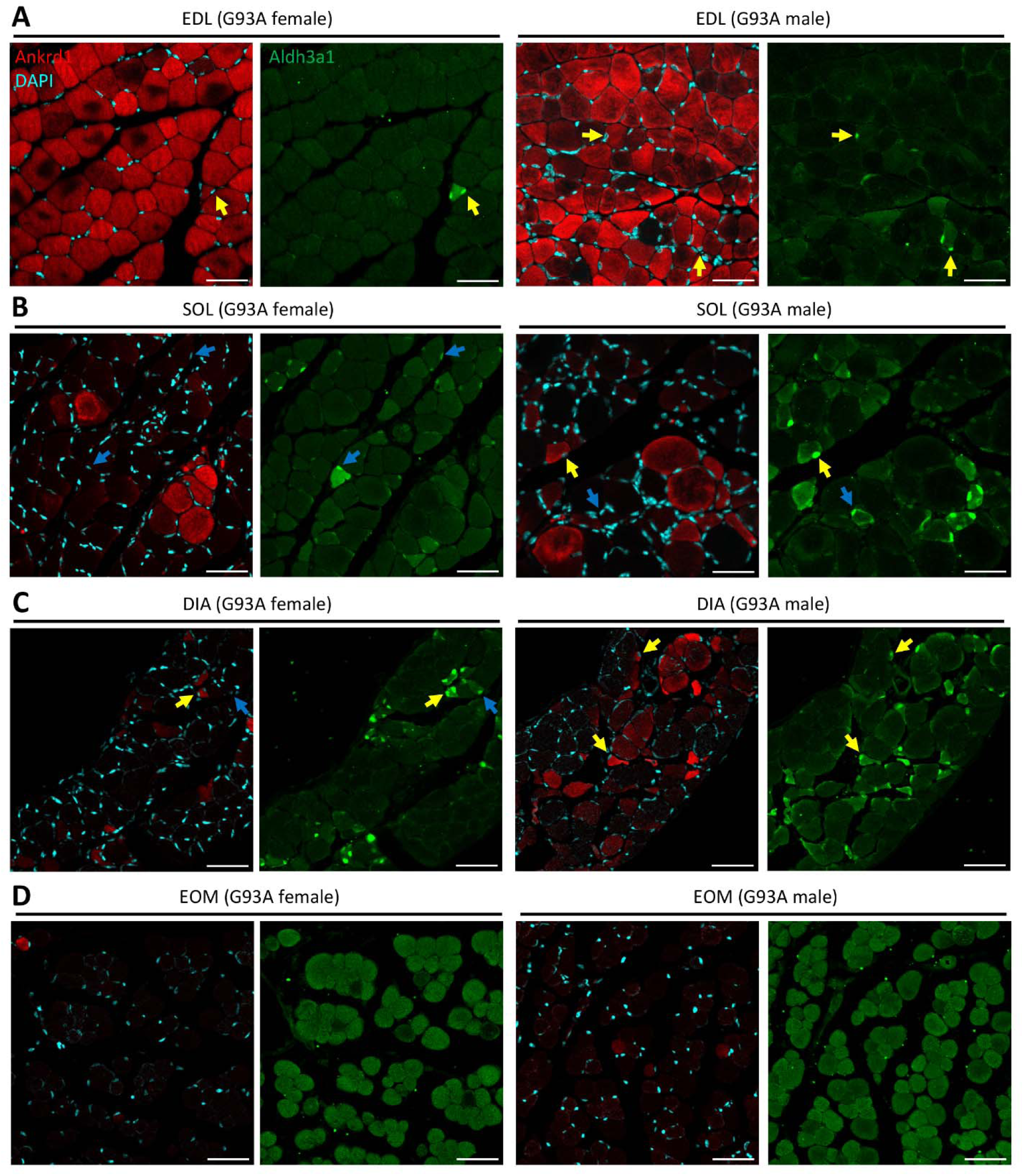
Section immunostaining results of Aldh3a1 and Ankrd1 in different muscles from end-stage G93A mice. (**A-D**) Transverse sections of EDL, soleus, diaphragm and EOMs from end-stage G93A mice stained with antibodies recognizing Aldh3a1 and Ankrd1. Yellow arrows denote nuclear Aldh3a1 positive myofibers that are also positive for Ankrd1. Blue arrows denote nuclear Aldh3a1 positive myofibers negative for Ankrd1. N = 6 (3 male and 3 female). Scale bars: 50 μm. Please also see **Figure 4-figure supplement 1** for section immunostaining results of Aldh3a1 in WT muscles.

### Sciatic nerve transection confirms the differential *Aldh3a1* and *Ankrd1* expression in EDL and soleus post denervation

Previous studies from us and others identified increased production of reactive oxygen species (ROS) in mitochondria as a common pathological feature in denervated muscles and muscle from ALS mouse models [38–40]. To better understand the relationship between *Aldh3a1* expression changes and muscle denervation, we performed sciatic nerve transection (SNT) in the right hindlimbs of WT mice 4-5 months of age. The left hindlimbs were sham operated as controls. The changes of gene expression in EDL and soleus with SNT (compared to their sham-operated controls) were examined at Day 3, 7 and 14 post operation by qRT-PCR (**Figure 5A** and **Figure 5-Source Data 1**). In EDL with SNT, there is no notable increase of *Aldh3a1* expression throughout the two weeks. In soleus with SNT, the elevated *Aldh3a1* expression was observed at Day 7 and 14. This time-dependent change of *Aldh3a1* expression was also confirmed at the protein level, with significant elevation detected in soleus but not in EDL post SNT (**Figure 5B**, **Figure 5-figure supplement 1** and **Figure 5-Source Data 2**). The *Ankrd1* expression was elevated in both EDL and soleus and peaked at Day 3 post SNT and decreased afterwards (**Figure 5**, **Figure 5-figure supplement 1** and **Figure 5-Source Data 3**). It is worth noticing that the increase of Ankrd1 protein was more pronounced in EDL (333-fold) compared to soleus (45-fold) at Day 3 post SNT, while the increase of Aldh3a1 protein was significant in soleus (7.6-fold) but not in EDL (1.6-fold) at Day 14 post SNT.

**Figure 5.**
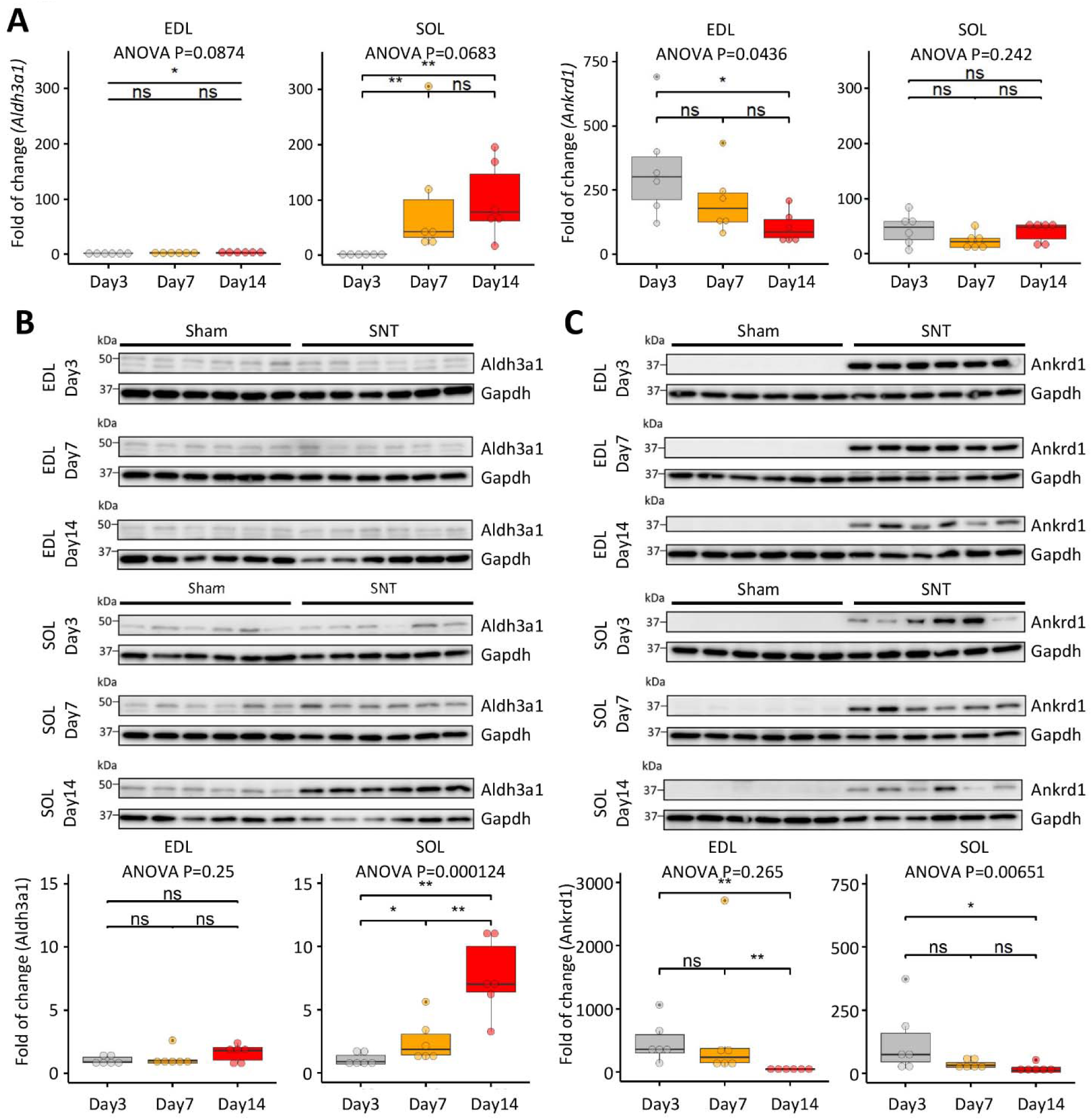
Time and muscle-dependent change of Aldh3a1 abundance after sciatic nerve transection. (**A**) Comparing the changes of *Aldh3a1* and *Ankrd1* expression levels in EDL and SOL from WT mice (4-5 months of age) collected 3 days, 7 days and 14 days after sciatic nerve transection (SNT). Fold of change refers to expression level differences between muscles with SNT and their sham-operated controls examined by qRT-PCR. N = 6 (3 male and 3 female). * P < 0.05; ** P < 0.01; ns, not significant (Wilcoxon rank-sum test). ANOVA P values are also shown. Please also see **Figure 5-Source Data 1**. (**B, C**) Comparing relative changes of Aldh3a1 and Ankrd1 protein (normalized by housekeeping protein Gapdh) in EDL and soleus (SOL) at 3 days, 7 days and 14 days after SNT to corresponding sham-operated controls by Western blot. N = 6 (3 male and 3 female). * P < 0.05; ** P < 0.01; ns, not significant (Wilcoxon rank-sum test). ANOVA P values are also shown. Please also see **Figure 5-figure supplement 1** and **Figure 5-Source Data 2, 3**.

The whole-mount immunostaining confirmed the complete denervation of myofibers in both EDL and soleus after SNT by the absence of axonal terminals in NMJs (**Figure 6A**). Myofibers with nuclear enriched Aldh3a1 was spotted in soleus at Day 7 post SNT. At Day 14, more myofibers showed increased Aldh3a1, not only in myonuclei, but also in cytosol (**Figure 6A**, insets). In line with the qPCR and Western Blot results, no noticeable increase of Aldh3a1 was observed in EDL myofibers after SNT compared to sham controls. Consistently, immunostaining revealed presence of Aldh3a1 positive myofibers in transverse sections of soleus but not EDL at Day 14 post SNT, whereas higher proportions of Ankrd1 positive myofibers were observed in these EDL muscles than soleus (**Figure 6B**). These data further demonstrate that *Aldh3a1* induction varies significantly between muscle types following denervation. Muscles with high proportions of mitochondria-rich oxidative fibers, like soleus and diaphragm, seem to be more prone to Aldh3a1 induction under elevated oxidative stress, likely as a self-protection mechanism.

**Figure 6.**
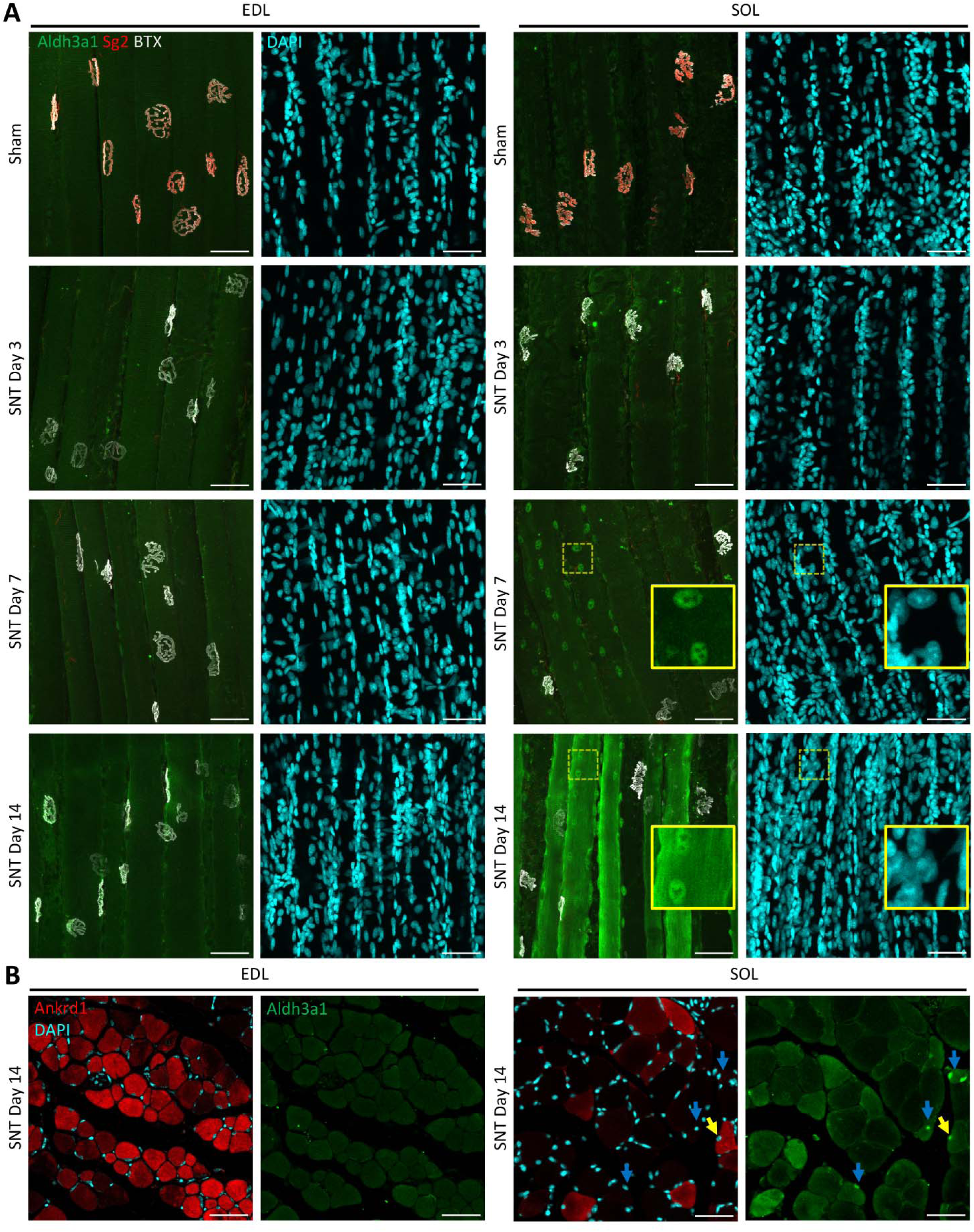
Examining subcellular distribution of Aldh3a1 protein in EDL and soleus muscles from WT mice with sciatic nerve transection. (**A**) Examining subcellular distribution of Aldh3a1 in EDL and soleus (SOL) muscles collected 3 days, 7 days and 14 days after SNT and sham-operated controls by whole-mount immunostaining. Antibodies against Sg2 labels axon terminals at NMJ and Alexa Fluor tagged BTX labels AChRs on the sarcolemma at NMJ. Regions denoted by dashed yellow boxes are enlarged in insets. N = 6 (3 male and 3 female). Scale bars: 50 μm. (**B**) Transverse sections of EDL and soleus muscles collected 14 days after SNT stained with Ankrd1, Aldh3a1 antibodies and DAPI. Yellow arrows denote nuclear Aldh3a1 positive myofibers that are also positive for Ankrd1. Blue arrows denote nuclear Aldh3a1 positive myofibers negative for Ankrd1. Scale bars: 50 μm.

### RNA-seq and pharmacological tests reveal potential mechanisms underlying muscle type dependent regulation of *Aldh3a1* expression

To explore the molecular mechanism underlying muscle type dependent regulation of *Aldh3a1* expression, whole muscle RNA-Seq were performed to compare between EDL, soleus, diaphragm and EOMs derived from end-stage G93A mice and age-matched WT controls (2 male and 2 female mice per group). In principle component analysis, EOM is the only muscle group with WT and G93A samples clustered together, whereas other muscle groups have more distinct expression profiles between WT and G93A (**Figure 7A**). This confirms that EOMs are less affected by ALS progression. To identify EOM signature genes, we generated lists of differentially expressed genes (DEGs, log2 fold change > 0.5, p.adjust < 0.05) comparing EOMs to EDL, soleus and diaphragm in WT mice. A total of 866 DEGs, including *Aldh3a1*, were in common between the three lists (**Figure 7-Source data 1**). Gene ontology analysis of these genes reveals that the most significant biological processes are related to eye development (**Figure 7-figure supplement 1A**). In addition, a good proportion of these genes are involved in axon guidance/neuron migration (**Figure 7-figure supplement 1B, C**), which is consistent with our previous discovery in EOM SC-derived myoblasts and myotubes [5] and may explain why neuromuscular junctions are preserved better in EOMs than other muscles in ALS.

**Figure 7.**
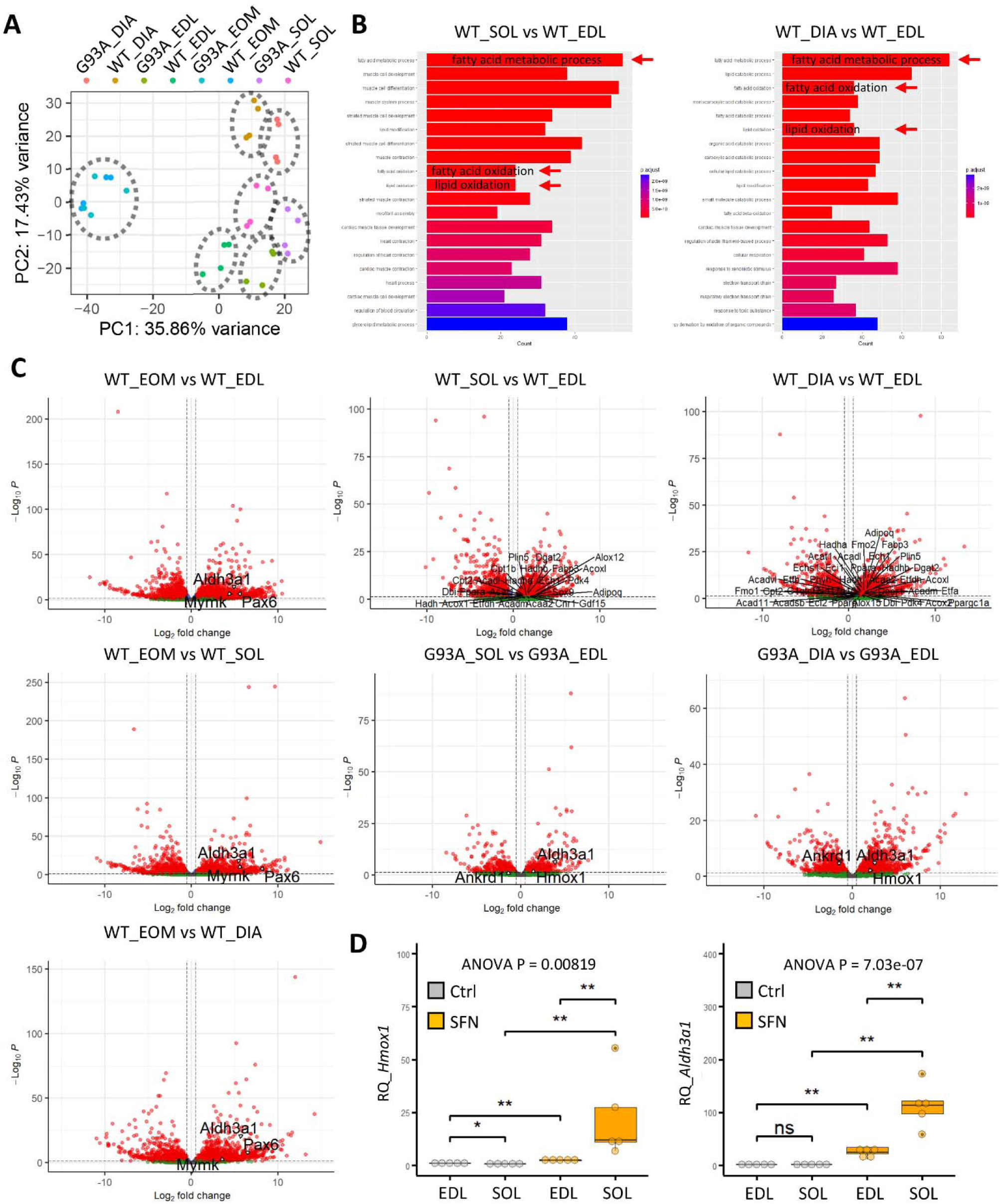
Whole muscle RNA-Seq reveals potential mechanisms underlying differential regulation of *Aldh3a1* expression between muscles. (**A**) Principle component analysis of different muscle samples from end-stage G93A mice and age-matched WT controls (2 male and 2 female mice per group). Dashed circles highlight samples with similar expression profiles. (**B**) Top 20 most significant gene ontology biological processes for differentially expressed genes (log2 fold change > 0.5, p.adjust < 0.05) comparing soleus and EDL muscles, diaphragm and EDL muscles in WT mice, respectively. Arrows highlight processes related to fatty acid metabolism and oxidation. (**C**) Volcano plots of genes differentially expressed between different muscle groups. The three plots in the left-most panel highlight three EOM signature genes *Aldh3a1*, *Mymk* and *Pax6* by arrowheads. In the upper two panels on the right, genes related to fatty acid/lipid oxidation in the gene ontology analysis above are denoted by lines. The lower two panels demonstrate the higher expression of *Aldh3a1*, *Hmox1* and the lower expression of *Ankrd1* in soleus and diaphragm compared to the EDL muscles from G93A mice (arrowheads). (D) qRT-PCR results of *Hmox1* and *Aldh3a1* in EDL and soleus SC-derived myotubes treated with DMSO (Ctrl) or 10 μM sulforaphane (SFN) for 48 hrs (N = 5 culture replicate). ** P < 0.01; * P < 0.05; ns, not significant (Wilcoxon rank-sum test). ANOVA P values are also shown. Please also see **Figure 7-source data 2**.

Since eye development is one of the top biological processes highlighted by gene ontology analysis, we suspected that the superior expression of *Aldh3a1* in EOMs was due to *Pax6*, one of the EOM signature genes (**Figure 7C**) that is a master regulator of eye formation and has been reported to activate *Aldh3a1* expression in mouse cornea [41]. However, upon closer inspection, we found that the transcripts per million (TPM) of *Pax6* is extremely low even for EOMs (the average is 1.47, not detected in one sample). Meanwhile *Pax6* transcripts were not detected in EOM SC-derived myoblasts or myotubes in our previous RNA-Seq results (GSE249484), which argues against our hypothesis. Another EOM signature gene that catches our attention is *Mymk* (**Figure 7C**), whose elevated expression in EOMs are also confirmed by qPCR (**Figure 1-figure supplement 1E**). *Mymk* mediates membrane fusion of myoblasts to existing myofibers during muscle development and regeneration [42]. Distinct from other muscles, EOMs are known for spontaneous activation of SCs to fuse with existing myofibers even without injury [5, 43]. That is why *Mymk* expression, which is supposed to be very low in uninjured WT muscles, is high in EOMs. Furthermore, our previous RNA-Seq data revealed notably higher expression of *Aldh3A1* in myoblasts than differentiated myotubes (GSE249484), implying that the spontaneous activation of SCs could contribute to the maintenance of high *Aldh3a1* levels in EOMs.

As to the differential regulation of *Aldh3a1* expression in other muscles, gene ontology analysis reveals that the most significantly upregulated biological processes in soleus and diaphragm compared to EDL in WT mice are fatty acid metabolism and oxidation (**Figure 7B**). Representative genes include *Acadl*, *Acadm*, *Acat1, Adipoq*, *Alox12* and *Alox15* (**Figure 7C** and **Figure 7-figure supplement 1D**). High basal level of lipid oxidation could predispose these muscles to stronger Nrf2 antioxidant response to additional oxidative stress caused by denervation or ALS [44]. Indeed, soleus and diaphragm showed a significant elevation of *Hmox1,* a classic target gene of Nrf2 signaling in G93A mice compared to EDL (**Figure 7C** and **Figure 1-figure supplement 1F**). Meanwhile more *Aldh3a1* and less *Ankrd1* transcripts were detected in soleus and diaphragm than EDL of G93A mice (**Figure 7C**), confirming the previous qPCR results.

Nrf2 signaling has been reported to activate *Aldh3a1* expression through the antioxidant response element in its promoter in human cancer cells [45, 46]. To examine whether Nrf2 dependent activation of *Aldh3a1* expression also occurs in skeletal muscle, we conducted pharmacological tests using Nrf2 signaling activator sulforaphane (SFN). The myotubes derived from EDL and soleus SCs (SCs were isolated from corresponding muscles in WT mice 4-5 months of age) were treated with 10 μM SFN for 48 hrs. As shown in **Figure 7D** and **Figure 7-Source Data 2**, more prominent upregulation of *Hmox1* and *Aldh3a1* expression were observed in soleus-SC derived myotubes than EDL-SC derived ones. These results support the differential predisposition to Nrf2 signaling activation as a contributor to differential regulation of *Aldh3a1* between muscles.

### Transducing myotubes with AAV-Aldh3a1 protects against 4-HNE induced apoptosis and plasma membrane defects

4-HNE is the most toxic reactive aldehyde generated during lipid peroxidation [47]. We evaluated 4-HNE levels in EDL and soleus muscles from end-stage G93A mice and age-matched WT controls through ELISA (**Figure 8A** and **Figure 8-Source Data 1**). 4-HNE levels were higher in soleus than EDL muscles from WT mice, potentially due to more active fatty acid oxidation in soleus as indicated in **Figure 7B**. However, 4-HNE level significantly increased in EDL but not in soleus muscles from end-stage G93A mice compared to WT controls, implicating the more pronounced induction of *Aldh3a1* expression in G93A soleus in preventing further elevation of 4-HNE levels under ALS.

**Figure 8.**
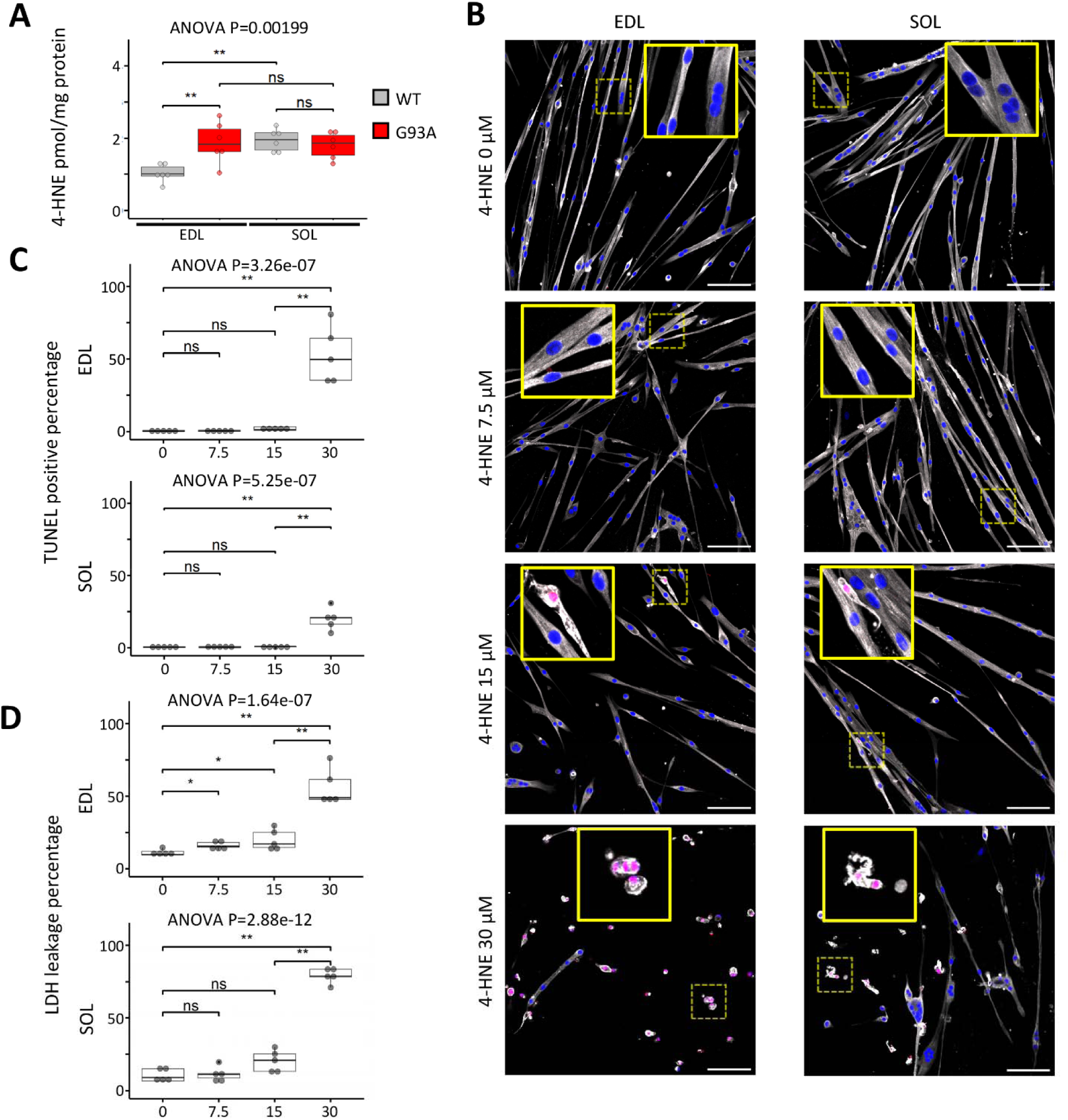
Characterizing 4-HNE cytotoxicity to myotubes derived from EDL and soleus satellite cells. (**A**) 4-HNE levels (normalized to protein amount) in EDL and soleus (SOL) muscles from end-stage G93A mice and age-matched WT controls (3 male and 3 female mice/group) measured by ELISA. ** P < 0.01; ns, not significant (Wilcoxon rank-sum test). ANOVA P value is also shown. Please also see **Figure 8-Source Data 1.** (**B**) Myotubes differentiated from EDL or soleus SCs isolated from 4-month-old WT mice for 5 days were treated with 0, 7.5, 15 or 30 μM 4-HNE for 2 hrs, respectively. Afterwards the myotubes were cultured for another 16 hrs in regular differentiation medium (as apoptosis takes time to occur) before fixation, TUNEL staining and immunostaining against MF-20 (labeling the cytosolic part of myotubes). DAPI counterstaining labelled the nuclei. N = 3 culture replicates. Regions denoted by dashed yellow boxes are enlarged in insets. Scale bars: 100 μm. (**C**) Averaged TUNEL positive percentage of nuclei in myotubes treated with different concentrations of 4-HNE (N = 5 culture replicates, 4 images analyzed and averaged for each culture replicate). ** P < 0.01; ns, not significant (t.test). ANOVA P values are also shown. Please also see **Figure 8-Source Data 2**. (**D**) Measuring 4-HNE cytotoxicity to myotubes using LDH leakage-based assay (N = 5 culture replicates). ** P < 0.01; * P < 0.05; ns, not significant (Wilcoxon rank-sum test). ANOVA P values are also shown. Please also see **Figure 8-Source Data 3**.

To assess 4-HNE cytotoxicity in skeletal muscles, the SCs-derived myotubes from WT EDL and soleus muscles were first treated with 0, 7.5, 15 or 30 μM of 4-HNE for 2 hours. Afterwards the myotubes were cultured for another 16 hours in differentiation medium before TUNEL staining, as apoptosis can take 12-24 hours to occur [48]. TUNEL positive nuclei (indicative of apoptotic cell death hallmarked by DNA fragmentation) were scarce in myotubes treated with 0, 7.5 or 15 μM 4-HNE but dramatically increased in myotubes treated with 30 μM 4-HNE (53% for myotubes derived from EDL myoblasts, 20% for myotubes derived from soleus myoblasts), accompanied by notable detachment of myotubes from the culture chamber (**Figure 8B, C** and **Figure 8-Source Data 2**). Due to the extensive occurrence of myotube detachment, calculating the percentage of TUNEL positive nuclei may not accurately reflect 4-HNE cytotoxicity. Since plasma membrane is commonly the primary site of 4-HNE production, we assessed the plasma membrane integrity by measuring the leakage of the cytosolic housekeeping enzyme lactate dehydrogenase (LDH) to the culture medium [49]. As shown in **Figure 8D** and **Figure 8-Source Data 3,** LDH leakage assay revealed a notable uptick of plasma membrane damage at 30 μM 4-HNE (65.6% for EDL-derived myotubes, 78.4% for soleus -derived myotubes).

Intriguingly, AAV-Aldh3a1 transduced myotubes, which had 2 to 6-fold higher levels of Aldh3a1 protein (**Figure 9A** and **Figure 9-Source Data 1**), exhibited dramatically lower percentage of apoptotic nuclei (3.5% for EDL-derived myotubes, 0.83% for soleus-dervied myotubes (**Figure 9B, C** and **Figure 9-Source Data 2**) and LDH leakage (30.4% for EDL-derived myotubes, 38.4% for soleus- derived myotubes (**Figure 9D** and **Figure 9-Source Data 3**). The transduction of AAV-Aldh3a1 alone did not change myotube viability compared to non-transduced controls (**Figure 9D**), indicating that Aldh3a1 protein is not toxic to cultured muscle cells.

**Figure 9.**
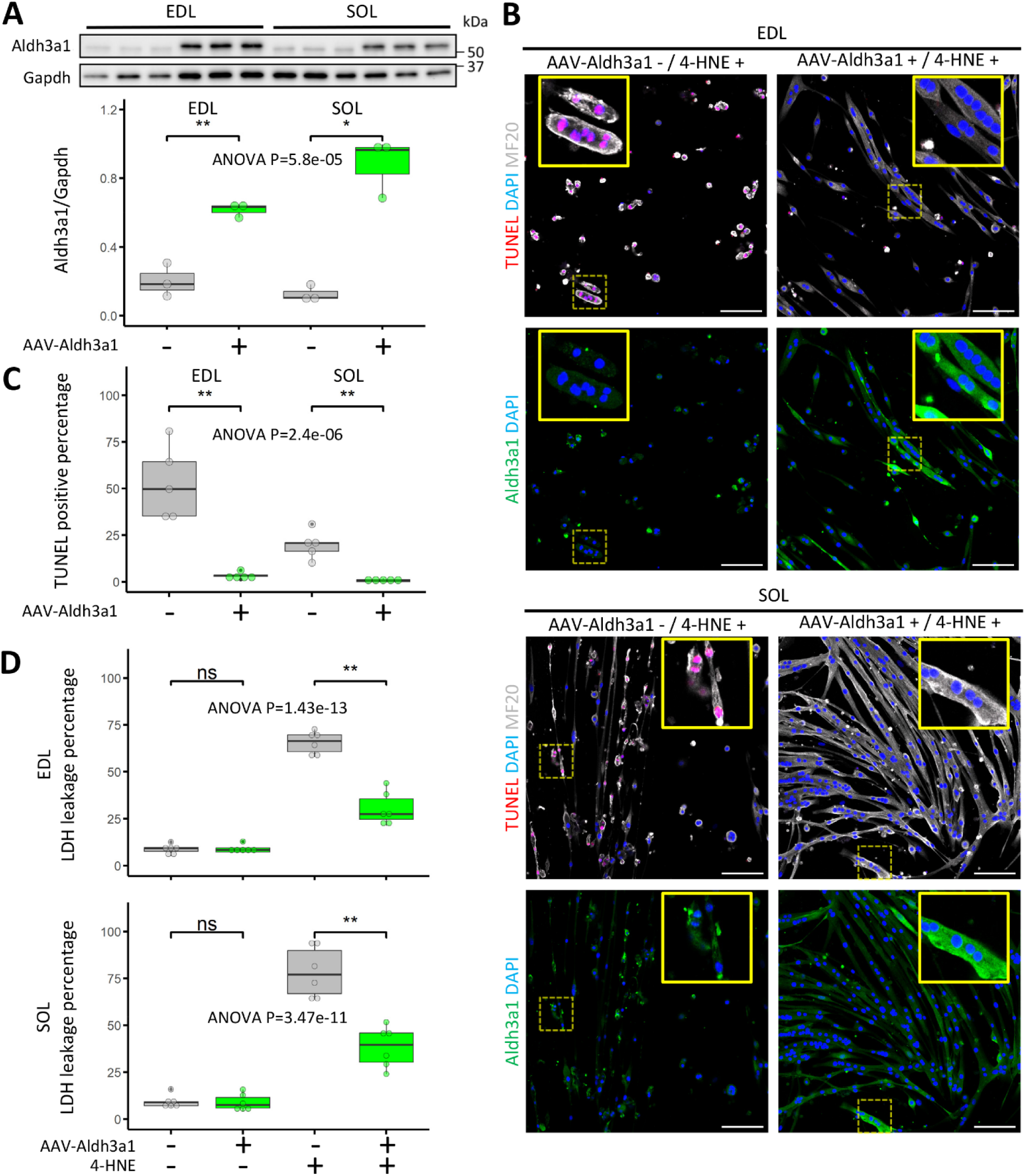
Transducing myotubes with AAV-Aldh3a1 protects against 4-HNE cytotoxicity. (**A**) Myotubes differentiated from EDL or soleus SCs for 1 day were incubated with or without AAV-Aldh3a1 for 2 days, followed by another 2-day culture in regular differentiation medium before protein extraction and Western blot. N = 3 culture replicates. ** P < 0.01; * P < 0.05 (t-test). Please also see **Figure 9-Source Data 1**. (**B**) Myotubes differentiated from EDL or soleus satellite cells isolated from 4-month-old female WT mice with or without AAV-Aldh3a1 transduction were treated with 30 μM 4-HNE for 2 hrs. Afterwards the myotubes were cultured for another 16 hrs in regular differentiation medium before fixation, TUNEL staining, immunostaining against MF-20 (labeling the cytosolic part of myotubes) and Aldh3a1. DAPI counterstaining labelled the nuclei. N = 3 culture replicates. Regions denoted by dashed yellow boxes are enlarged in insets. Scale bars: 100 μm. (**C**) Quantification results of averaged TUNEL positive percentage of nuclei in myotubes (N = 3 culture replicates, 4 images analyzed and averaged for each culture replicate). ANOVA P values are also shown. * P < 0.05 (t-test). Please also see **Figure 9-Source Data 2**. (**D**) Measuring the protective effect of AAV-Aldh3a1 against 4-HNE cytotoxicity to myotubes using LDH leakage-based assay (N = 6 culture replicates). ** P < 0.01; ns, not significant (Wilcoxon rank-sum test). ANOVA P values are also shown. Please also see **Figure 9-Source Data 3**.

Increased LDH leakage and myotube detachment following 4-HNE treatment indicate compromised plasma membrane integrity. We previously reported defects in MG53 mediated membrane repair mechanism G93A mice [35, 50]. Here we examined the impact of 4-HNE treatment on the formation of stable MG53 repair patches in myotubes with overexpression of GFP-MG53 [34]. Without 4-HNE treatment, GFP-MG53 was predominantly cytosolic and quickly translocated to the plasma membrane in response to cell membrane injury induced by saponin, a compound permeabilizing plasma membrane. About 86% recorded myotubes maintained elevated membranous GFP-MG53 signal at the end of recording, implying the formation of stable MG53 repair patches (**Video 1, Figure 10** and **Figure 10-Source Data 1**), while the proportion dropped to 8.6% for myotubes treated with 30 μM 4-HNE for 2 hours before recording (**Figure 10B, D** and **Figure 10-Source Data 1**). A good portion of 4-HNE treated myotubes already had blebs on the plasma membrane before saponin administration. Additionally, the peak intensity of membranous GFP-MG53 signals were also lower than those not treated with 4-HNE (**Figure 10C**). These observations indicate that plasma membrane damage has already occurred during 4-HNE treatment (**Video 2** and **Figure 10**). Intriguingly, pretreatment of myotubes with AAV-Aldh3a1 before 4-HNE treatment not only prevented 4-HNE induced plasma membrane blebbing, restored the peak intensity of membranous GFP-MG53 signal, but also increased the proportion of myotubes maintaining elevated membranous GFP-MG53 signal to 77% (**Video 3**, **Figure 10** and **Figure 10-Source Data 1**).

**Figure 10.**
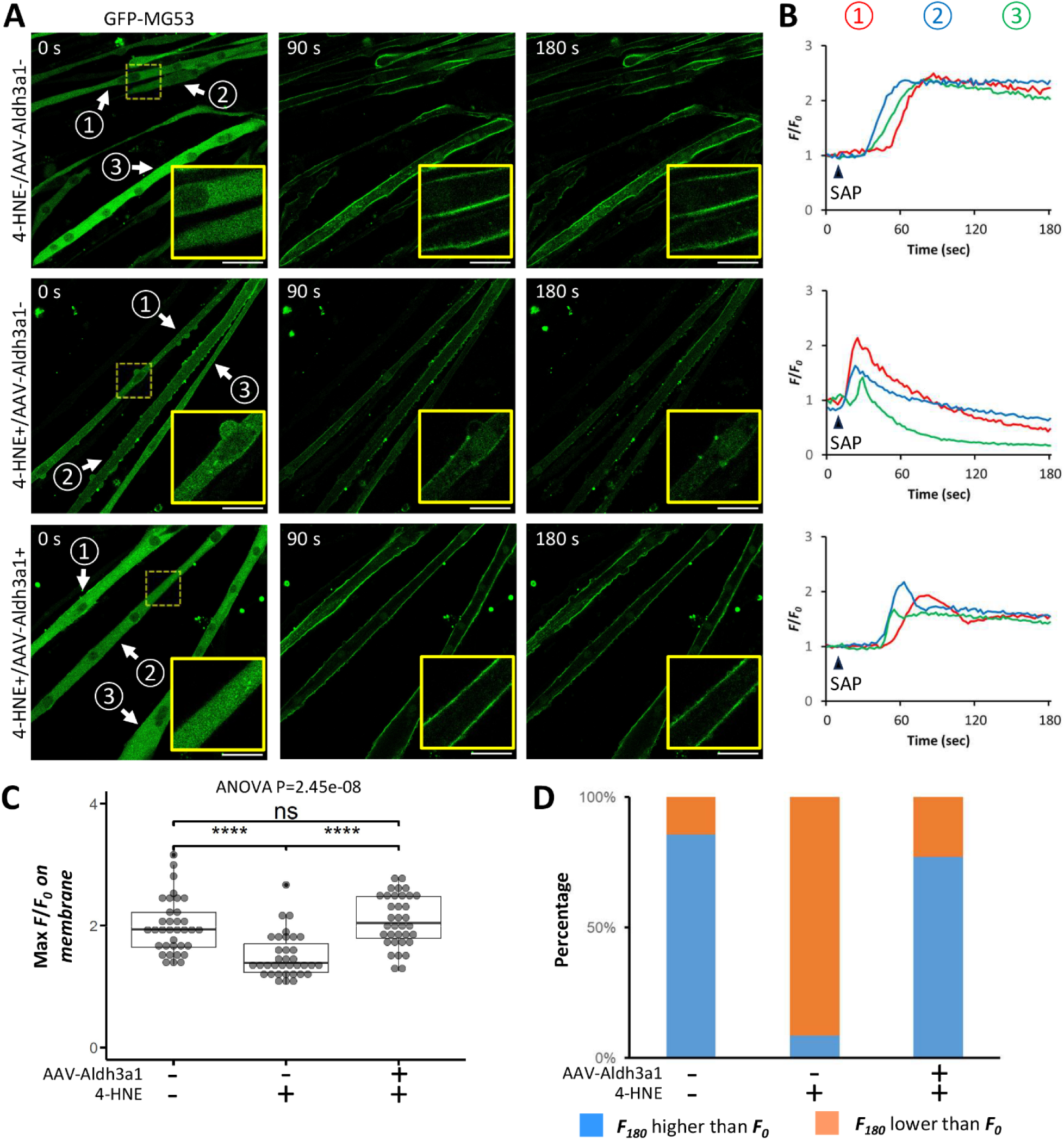
AAV-Aldh3a1 protects against 4-HNE compromised stabilization of MG53 repair patches in partially permeabilized myotubes. (**A**) Time lapse imaging of saponin-induced formation of GFP-MG53 repair patches on the plasma membrane in soleus SC-derived myotubes with or without AAV-Aldh3a1 transduction and/or 4-HNE treatment (30 μM for 2 hrs). Regions denoted by dashed yellow boxes are enlarged in insets. Arrows highlight myotubes whose GFP- MG53 intensity on the plasma membrane were profiled in panel B. Scale bars: 50 μm. Please also see Videos 1-3. (**B**) Plotting background-corrected, initial timepoint intensity normalized signal of GFP-MG53 on plasma membrane (*F*/*F_0_*) over time for the three myotubes indicated in Panel A. Arrowheads denote the time point when saponin was applied. (**C**) Maximum *F*/*F_0_*of recorded myotubes under different treatment conditions (N = 35 for each group). **** P < 0.0001; ns, not significant (Wilcoxon rank-sum test). ANOVA P values are also shown. Please also see **Figure 10-Source Data 1**. (**D**) Percentage stacked column chart highlights the proportion of recorded myotubes with *F_180_* (end of recording) higher than *F_0_* (blue) and the proportion of those with *F_180_* lower than *F_0_* (orange) in different treatment groups. Please also see **Figure 10-source data 1**.

## Discussion

In this study we discovered that Aldh3a1 protein is extremely abundant in mouse EOMs and can be upregulated in other muscles under pathological conditions such as denervation and ALS progression. The extent of *Aldh3a1* upregulation in the ALS mouse model (G93A) varies over muscle type, more prominent in soleus, diaphragm than EDL. In contrast, the muscle pathological remodeling marker *Ankrd1* is more substantially elevated in EDL than soleus and diaphragm of G93A mice. The induction pattern of *Aldh3a1* is also inverse to that of *Ankrd1* over muscle type and time in WT mice with sciatic nerve transection. In addition, the protective effects of Aldh3a1 against 4-HNE in muscle are multi-faceted, from DNA fragmentation in the nuclei to plasma membrane damage and repair defects involving MG53.

Aldh3a1 has been reported to be an inactivation-resistant detoxifier of reactive aldehydes such as 4-HNE in an enzyme kinetics study [27]. Here we detected increased 4-NHE levels in G93A muscles without pronounced increase of Aldh3a1. Meanwhile 4-HNE treated myotubes showed an increase in nuclear DNA fragmentation, cell membrane leakage and loss of MG53 membrane repair function, implying that elevated 4-HNE level overwhelmed the physiological detoxification capacity of muscle cells. Remarkably, pretreating myotubes with AAV-Aldh3a1 potently protected myotubes against all the above cytotoxic effects caused by 4-HNE. In addition, AAV-Aldh3a1 transduction alone did not alter myotube viability, which increases the likelihood for its therapeutic application in the future.

Nuclear enrichment of Aldh3a1 has been observed previously in cultured human and rabbit corneal epithelial cells overexpressing *Aldh3a1* [29, 51, 52]. Since reactive aldehydes such as 4-HNE can directly modify genomic DNA, leading to mutagenesis and fragmentation in multiple non-muscle cell types [16, 21, 22], the DNA protection role of Aldh3a1 has been investigated. Indeed, it was reported to prevent DNA damage and apoptosis induced by 4-HNE, hydrogen peroxide, or UVR in cultured corneal epithelial (HCE) cells and stromal fibroblasts [20, 51–54]. Aside from its role to directly detoxify reactive aldehydes, Aldh3a1 was also reported to promote a series of DNA damage detection/repair related processes in HCE cells including prolonging cell cycle, increasing levels of total and phospho (Ser15) p53 and activating ATM/ATR signaling pathway, which is central to the maintenance of genome integrity [51, 52, 54–56]. These effects could be a unique advantage of applying Aldh3a1 in therapy. Interestingly, in HCE cells overexpressing *Aldh3a1,* the expression of *GADD45A*, another muscle denervation marker [57], was found to be downregulated [54], indicating the inverse correlation between the expression levels of *Aldh3a1* and muscle denervation marker *Ankrd1* may not be a coincidence, but a phenomenon broadly presents in different tissues. In this study, we did kymographic profiling of Aldh3a1 inside myonuclei and unveiled its preferential distribution in the euchromatin regions, where genomic DNA is less compacted and could be more vulnerable to attacks by reactive aldehydes. It is reasonable to hypothesize that *Aldh3a1* upregulation in muscle is a self-defense mechanism against pathologically elevated lipid peroxidation stress not only to protect cytosolic protein but also to maintain genome DNA integrity.

An alternative scenario would be Aldh3a1 self-regulates its own expression via a positive feedback loop. Yet this is unlikely as Aldh3a1 is not known to directly bind DNA like a transcription factor. It is also worth noticing that the high expression of *Aldh3a1* gene does not guarantee the nuclear enrichment of Aldh3a1 protein. In EOMs and myotubes transduced with AAV-Aldh3a1, we did not observe notable nuclear enrichment of Aldh3a1 (**Figure 3A**, **Figure 3-figure supplement 1A**, **Figure 4D**, **Figure 4-figure supplement 1** and **Figure 9B**). The mechanism driving Aldh3a1 nuclear translocation is still under investigation, as the deactivation of its predicted nuclear localization sequence did not achieve the expected outcome [52]. It is possible that the extent of Aldh3a1 unclear accumulation depends on the status of the oxidative stress inside the nuclei.

*Aldh3a1* expression has been reported to be activated by Nrf2 signaling through antioxidant response element (ARE) in its promoter in human cancer cells [45]. The activation of Nrf2 signaling has been reported to enhance exercise endurance capacity, augment skeletal muscle regeneration after ischemia-reperfusion injury and ameliorate muscle mass/contractility decline during aging [58–63]. In addition, Nrf2 signaling activator edaravone (Radicava) has been approved by FDA to combat ALS [64, 65].

Our whole muscle RNA-Seq detected higher expression levels of lipid oxidation related genes in soleus and diaphragm than EDL muscles. This could be due to higher content of oxidative myofibers in these two muscles [66], whose mitochondria have a predisposition toward using fatty acids as fuel [67]. This is also supported by higher 4-HNE levels in soleus than EDL muscles from WT mice indicated by ELISA. The high basal lipid oxidation levels could predispose soleus and diaphragm to Nrf2 signaling activation [44], potentially through making the promoter regions of antioxidant genes more accessible to Nrf2. In addition, the higher proportion of mitochondria-rich oxidative fibers in soleus and diaphragm could also accelerate the activation of Nrf2 signaling as a self-defense mechanism due to faster accumulation of mitochondria-produced ROS under ALS or denervation [38–40]. Indeed, we detected more transcripts of Nrf2 target gene *Hmox1* in soleus and diaphragm than EDL of G93A mice. In addition, treatment of Nrf2 activator SFN induced more prominent upregulation of *Hmox1* and *Aldh3a1* in soleus-SC derived myotubes than EDL-SC derived ones. Thus, the differential predisposition to Nrf2 signaling activation is likely a contributing factor to the differential regulation of *Aldh3a1* expression between soleus, diaphragm and EDL muscles.

However, Nrf2 signaling alone cannot explain the superior expression of *Aldh3a1* in EOMs as none of its classic target genes, such as *Hmox1*, *Nqo1*, *gclm* or *gsta4* [64], shows up in the EOM signature gene list. Based on the results of gene ontology analysis, we suspected that eye development-related genes, such as *Pax6*, were involved in activating *Aldh3a1* expression in EOMs [41]. However, the extremely low levels of *Pax6* transcripts in EOM samples, combined with their absence in EOM-SC derived myoblasts and myotubes in our previous RNA-Seq data (GSE249484), argues that they are more likely due to residual eye tissue left during dissection, rather than endogenous expression in the muscles. On the other hand, the identification of *Mymk* as an EOM signature gene provides an alternative scenario to explain high *Aldh3a1* levels in EOMs. *Mymk* is known for mediating the fusion of myoblasts with existing myofibers during muscle regeneration [42]. Its expression should be low in uninjured muscles, but EOM is an exception because of spontaneous activation of SCs. This unique property was first documented due to the observation of BrdU positive nuclei (from proliferating myoblasts) in the center of uninjured EOM myofibers [43]. Our recent fluorescence-activated cell sorting analysis also revealed significantly more heterogenous levels of VCAM (quiescent satellite cell marker) in EOM SCs than those from other muscles of WT mice, confirming spontaneous activation of EOM SCs without pathological stimuli [5]. Furthermore, we detected notably more *Aldh3a1* transcripts in myoblasts than myotubes (either originated from EOM, diaphragm or hindlimb muscle SCs) in previous RNA-Seq results (GSE249484). This is not surprising as many types of stem cells are characterized by upregulated expression of specific ALDH isoenzymes, like *Aldh1a1*, *Aldh1a3* and *Aldh3a1*, which promote their survival [68–70]. High *Aldh3a1* expression has been correlated with cell proliferation and resistance to reactive aldehyde cytotoxicity [70]. Therefore, the spontaneous activation of abundant SCs in EOMs and their fusion with existing myofibers could contribute to high *Aldh3a1* levels.

As to 4-HNE targets in ALS patients, a few key proteins in the ventral horn motor neurons, such as the astrocytic glutamate transporter EAAT2, have been found to be modified by 4-HNE [71]. In ALS rodent models, 4-HNE has been reported to form adducts with Drp2, Hsp70, glutathione and carnosine [72, 73]. Here, we demonstrated functional disruption of MG53 by 4-HNE. Previous studies implied cysteine 242 of mouse MG53 as a redox sensor to oligomerize or crosslink [34, 74]. Meanwhile cysteine is the residue most prone to 4-HNE attack through Michael addition [75, 76]. Interestingly, similar to the time-lapse imaging results of 4-HNE treated myotubes expressing GFP- MG53 we reported here, cardiomyocytes expressing GFP-MG53(C242A), which mutates cysteine 242 to alanine, also demonstrated failed maintenance of membranous signal after the application of saponin [74]. Thus, whether C242 of MG53 is one of the residues forming Michael adducts with 4- HNE in muscles of G93A mice is worth further investigation in future studies. The result may explain the abnormal formation of cytosolic MG53 aggregates in these muscles [35].

In summary, we identified multi-faceted benefits of Aldh3a1 in protecting against reactive aldehyde cytotoxicity, which likely contribute to the distinct muscle response observed in ALS progression. The spectrum of vulnerability among motor neurons innervating different muscles remains a mystery in ALS. Specifically, the ocular motor neurons are notably resistant to ALS, persisting throughout the disease [77]. While our results do not exclude a primary effect of those neurons on EOMs, we can only speculate that ocular motor neurons, like EOMs, may possess a superior capacity to mitigate lipid peroxidation cytotoxicity due to their close developmental relationship with the eye. Future studies are warranted to experimentally validate this hypothesis and test the therapeutic potential of motor neuron targeted expression of Aldh3a1 for ALS.

## Materials and methods

### Key Resources Table

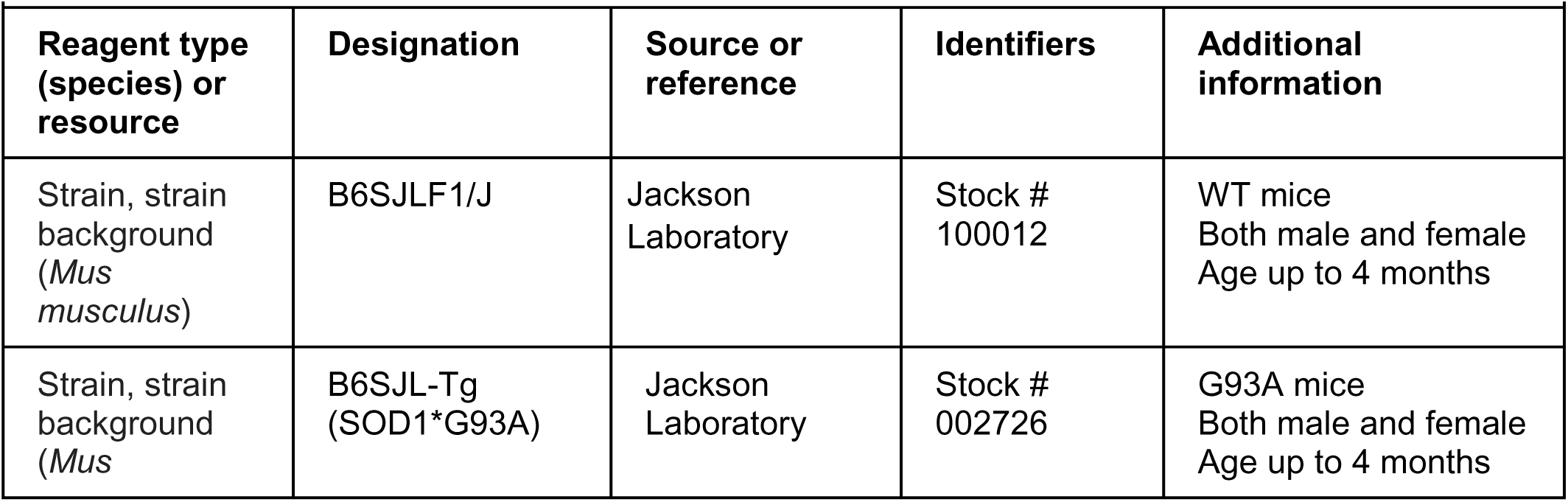
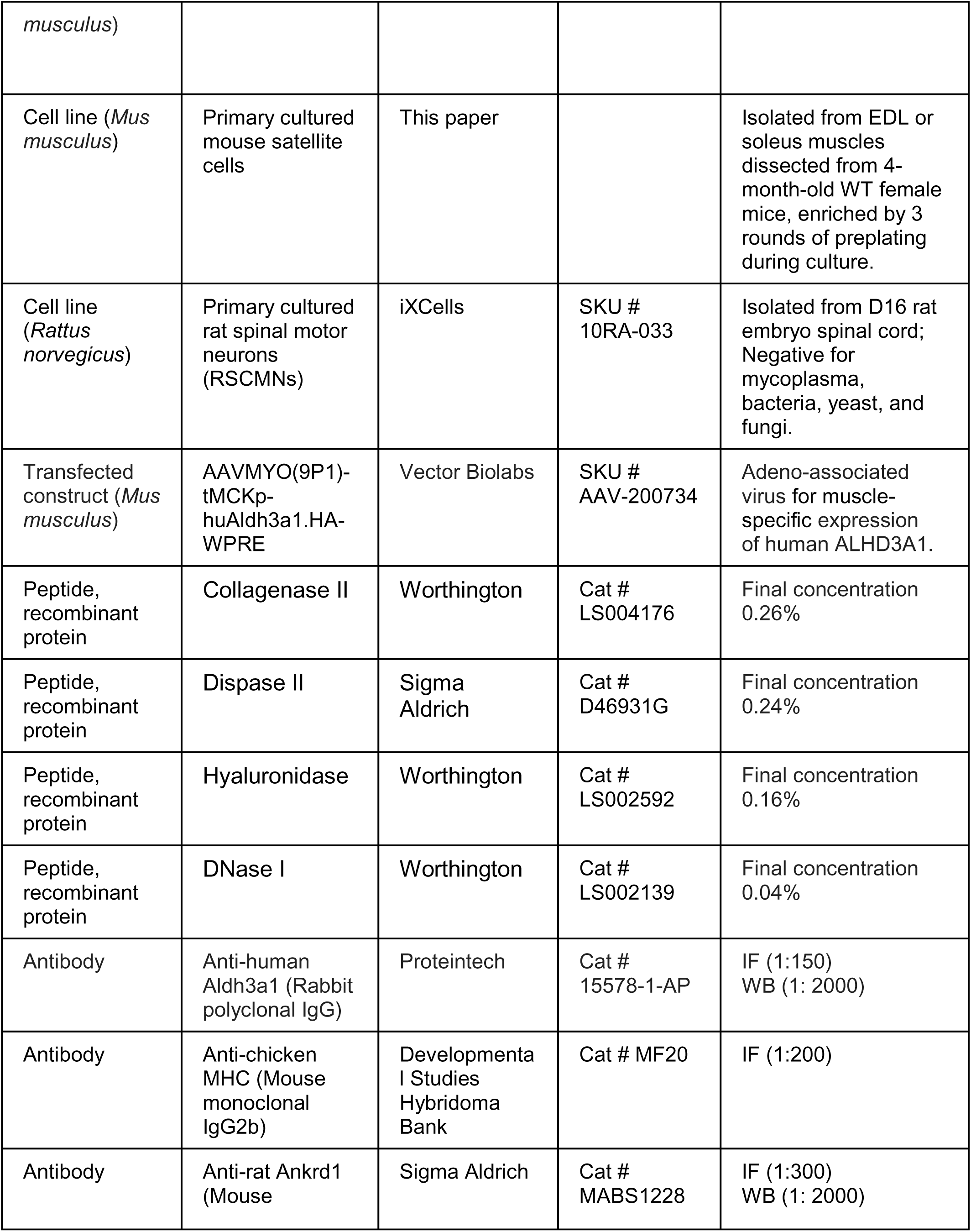
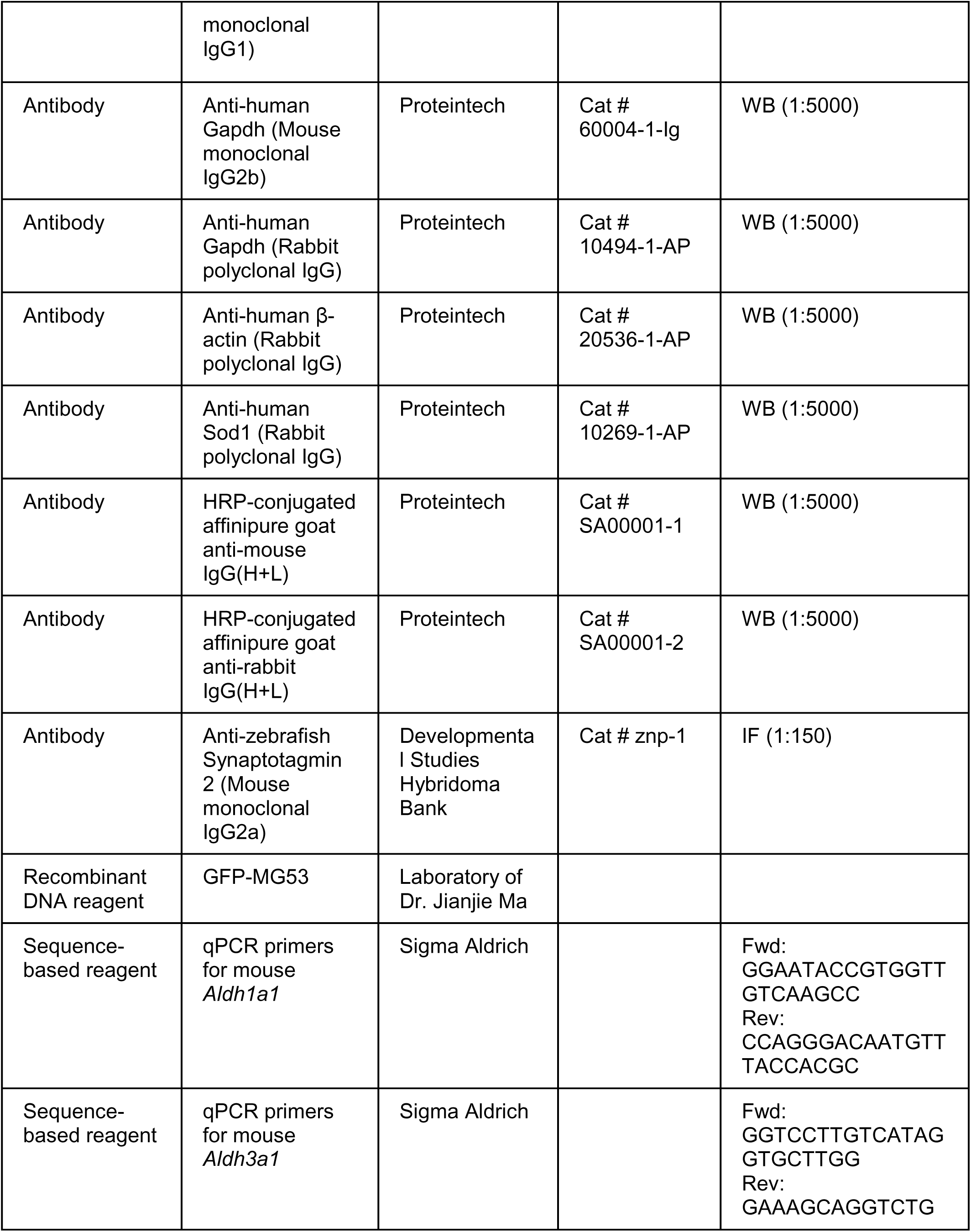
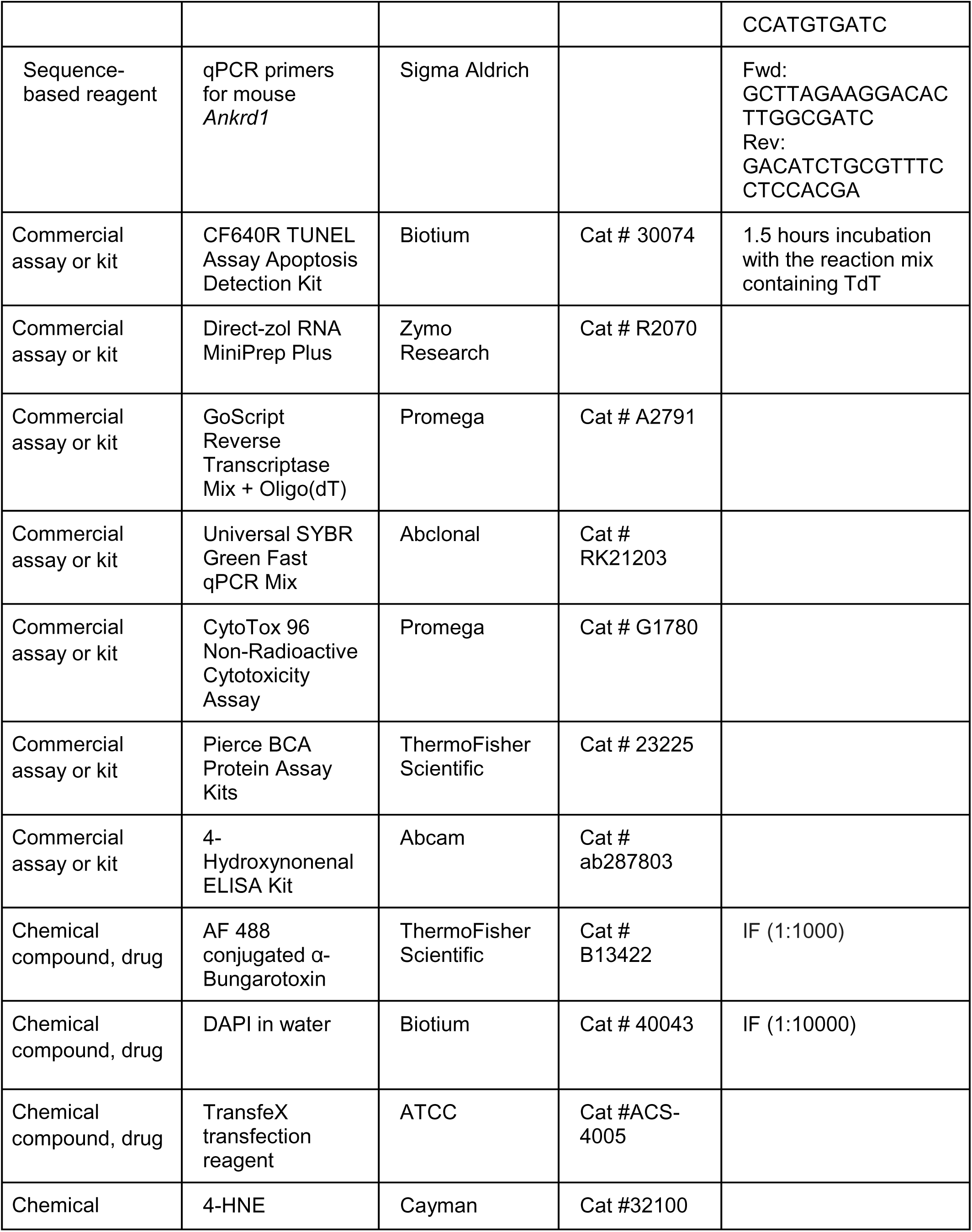
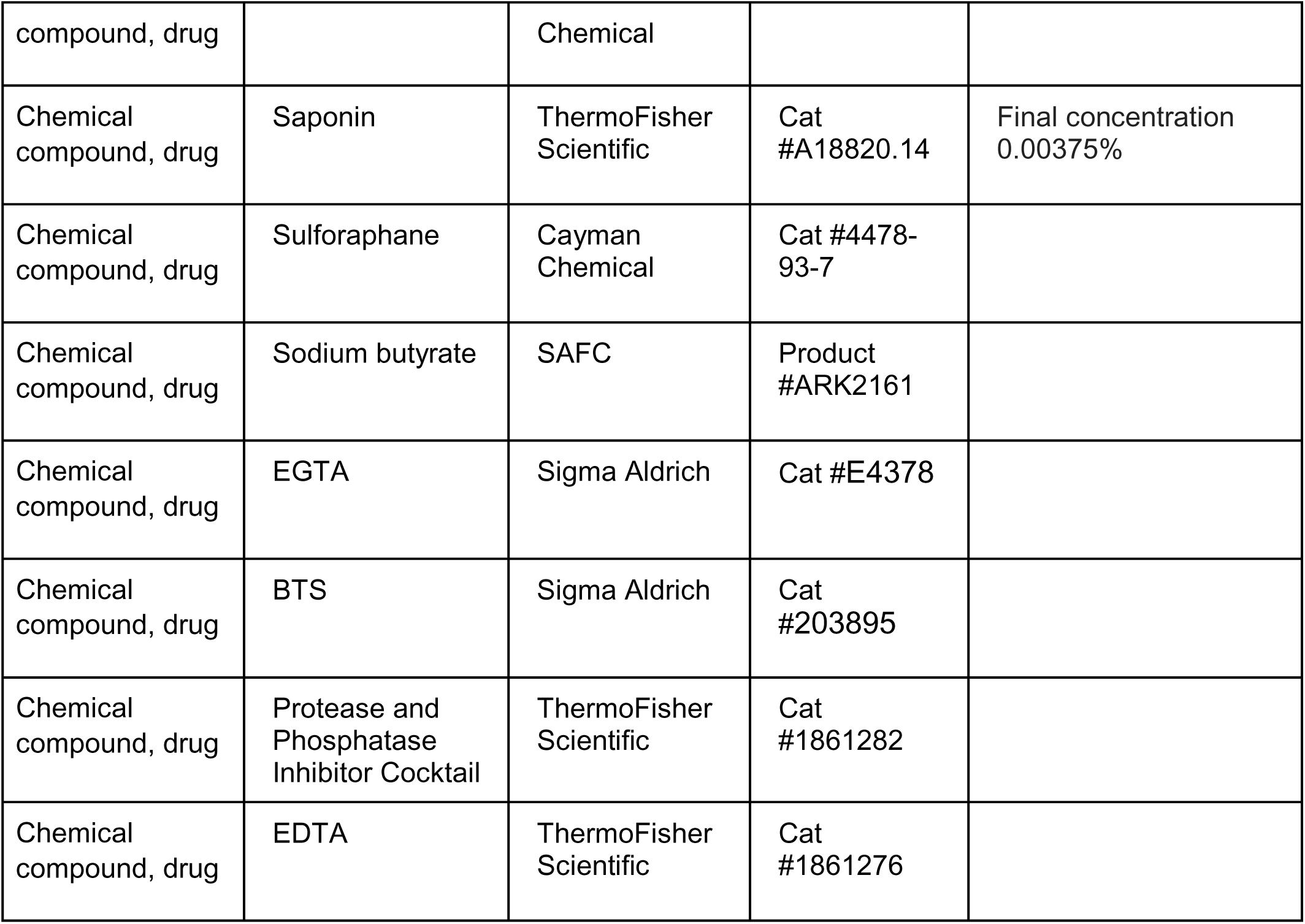

### Animals

All animal experiments were carried out in accordance with the recommendations in the *Guide for the Care and Use of Laboratory Animals* of the National Institutes of Health. The protocol on the usage of mice was approved by the Institutional Animal Care and Use Committee of the University of Texas at Arlington (A19.001, approval date: 09/20/2018). WT mice used in this study were of B6SJL background. The ALS transgenic mouse model (hSOD1G93A) with genetic background of B6SJL was originally generated by Drs. Deng and Siddique’s group at Northwestern University and deposited to the Jackson Lab as B6SJL-Tg (SOD1*G93A) [31]. G93A mice of both genders were euthanized for sample collection at the endpoint when they were unable to right themselves to a sternal position within 30 seconds (4-5 months of age). WT mice of the corresponding gender were euthanized for sample collection at the same day as their G93A littermates.

### Sciatic nerve transection

Surgical instruments were sterilized by autoclave before operation. Mice were anesthetized with constant flow isoflurane inhalation. Hairs in the two posterior thighs and lower back were shaved as much as possible with electric clipper. The skin region to be operated on was aseptically prepared using surgical scrub with betadine or equivalent surgical soap and rinse with 70% alcohol. Incision through the skin and superficial muscles was made parallel and just inferior to the femur of the right hindlimb. Curved-end forceps were used to divide the muscles and expose the sciatic nerve. 5 mm of the sciatic nerve was removed with fine surgical scissors. Both ends of the nerve were sutured to prevent regeneration. The incision on the skin was closed with stainless steel wound clips. A sham procedure following the same steps without severing the sciatic nerve was performed for the left hindlimb as control.

### Immunofluorescence (IF) and imaging of whole mount and muscle samples

For whole mount immunofluorescence, EDL, soleus, diaphragm and EOM samples were fixed and permeabilized in precooled methanol at -20 °C for 15 min. The samples were rehydrated by three changes of PBS and incubated with Alexa Fluor 488 conjugated α-Bungarotoxin (ThermoFisher B13422, 1:1000) in blocking buffer (PBS containing 2% BSA, 2% horse serum, 0.1% Tween-20, 0.1% Triton X-100 and 0.05% sodium azide) at 4 °C for 1 day. The samples were then washed with PBS and further incubated with the primary antibodies against Aldh3a1 (Proteintech 15578-1-AP 1:150) and Synaptotagmin-2 (Sg2 for short, DSHB ZNP-1 concentrate + 50% glycerol 1:150) in blocking buffer at 4 °C for 1 day. On the third day, the samples were washed with PBS for 3 times and incubated with corresponding secondary antibodies labelled with Alexa Fluor.

For section immunofluorescence, EDL, soleus, diaphragm and EOM were fixed in either 4% paraformaldehyde or 3% glyoxal fixative (pH 4.5, containing 20% ethanol) for 8-12 hours at 4 °C. Glyoxal based fixative can penetrate tissue faster due to the presence of ethanol [78–80]. Antigen retrieval was conducted in Tris-EDTA buffer (pH 9.0) for 30 min at 95 °C. Afterwards the samples were washed with PBS containing 1% glycine once and two more times with PBS. Then the samples were immersed in blocking buffer for 45 min at room temperature followed by primary antibody incubation at 4 °C overnight. Next day, after washing with PBS for three times, the samples were incubated with Alexa Fluor labelled secondary antibodies (ThermoFisher Scientific 1:800) for 2 hours at room temperature. The samples were then washed with PBS, counterstained with DAPI and mounted in antifade mounting media (20 mM Tris, 0.5% N-propyl gallate, 80% glycerol) for imaging. Primary antibodies used: Aldh3a1, Ankrd1 (Sigma MABS1228 1:200)

### RNA extraction and qRT-PCR

Homogenization was performed in FastPrep-24 Classic bead beating grinder (MP Biomedicals 116004500). The muscle homogenate was transferred to centrifuge tubes containing phase lock gel (QuantaBio 2302830) and 1/10 volume of 1- bromo-3-chloropropane was added. The tubes were hand shaken for 12 seconds, left at bench top for 2 minutes and then centrifuged at 16000 x g at 4 °C for 15 minutes. The upper phase was transferred to a new centrifuge tube and mixed with equal volume of ethanol. The following steps of RNA purification were performed with Direct-zol RNA Miniprep Plus kit (Zymo Research R2070). RNA concentration was measured with Quantus Fluorometer (Promega E6150). GoScript Reverse Transcription Mix, oligo(dT) was used for the reverse transcription reaction (Promega A2791). First-strand cDNAs were diluted and mixed with 2X Universal SYBR Green Fast qPCR Mix (Abclonal RK21203), as well as corresponding primers for qPCR using StepOnePlus Real-Time PCR system (ThermoFisher Scientific 4376600). Relative quantification (RQ) of gene expression was generated by ΔΔCt method. The sequences of primers used for qPCR were listed in Key Resource Table.

### Bulk RNA-Seq and analysis

RNAs were extracted from EDL, soleus, diaphragm and extraocular muscles from end-stage G93A mice and age-matched WT controls (2 male and 2 female mice per group) as described above. cDNA library preparation and sequencing were carried out by Innomics.inc. The average number of clean reads per sample was 23 million. The clean reads were mapped to Genome Reference Consortium Mouse Build 39 (GRCm39) using Rsubread package of Bioconductor [81] and the average percentage of aligned reads was 87%. DESeq2 package [82] was used to identify differentially expressed genes (DEGs) between different samples. The data were prefiltered to keep only genes with counts ≥ 10 in at least 4 samples. Only genes with log2FC > 0.5 and p.adjust < 0.05 were considered differentially expressed in volcano plots and gene ontology analysis. Principle component analysis (PCA) was performed with pcaExplorer package [83, 84]. Volcano plots were created using EnhancedVolcano package. Gene ontology analysis was performed with clusterProfiler package [85]. Fastq files of clean reads and the excel file containing TPM values will be uploaded to GEO database upon acceptance of the manuscript.

### Western blot

Proteins were extracted by grinding corresponding muscles in 10 volumes of RIPA buffer containing protease inhibitors (ThermoFisher Scientific), resolved by SDS-PAGE, and transferred to PVDF membrane with Bio-Rad semidry transfer cell. The primary antibodies were Aldh3a1 (Proteintech 15578-1-AP), Ankrd1 (Sigma MABS1228) and Gapdh (Proteintech 60004-1-Ig). Protein bands were detected with Bio-Rad Clarity ECL kit and ChemiDoc Imaging system. Signal strengths and backgrounds were measured using ImageJ software.

### BCA protein assay and 4-HNE enzyme-linked immunosorbent assay (ELISA)

To evaluate 4-HNE levels in EDL and soleus muscles from end-stage G93A mice and age- matched WT controls. We homogenized corresponding muscles in lysis buffer (50mM Tris + 0.9% NaCl + 0.1%SDS + Protease and Phosphatase Inhibitor Cocktail + 1 mM EDTA) using FastPrep-24 Classic bead beating grinder (MP Biomedicals 116004500). The homogenates were briefly centrifuged and transferred to new 1.5 ml microcentrifuge tubes. The tubes were centrifuged again for 15 min at 12000 x g and the supernatants were collected for concentration measurement using Pierce BCA Protein Assay Kit (ThermoFisher Scientific 23225) according to their protocol on a plate reader (SpectraMax i3x, Molecular Devices).

The homogenates were then diluted to ∼1 mg/ml with the sample/standard dilution buffer of 4- HNE ELISA kit (Abcam ab287803). Each well in the Micro ELISA Plate of the kit was washed twice with 1x wash solution. 50 μl of diluted samples or 4-HNE standards were loaded to each well of the Micro ELISA Plate and incubated with 50 μl of diluted biotin-detection antibodies at 37 °C for 45 min. The plate was washed again with 1x wash solution for 3 times and incubated with 0.1 ml SABC working solution/well at 37 °C for 30 min. The plate was then washed again with 1x wash solution for 5 times and loaded with 90 μl TMB substrate solution/well at 37 °C for 15 min. 50 μl stop solution was added per well and the absorbance of each well at 450 nm and 620 nm were immediately measured by a plate reader. 4-HNE concentration derived from the standard curve was normalized to the protein concentration of the corresponding well.

### Primary culture of myoblasts

EDL or soleus muscles were dissected from two 4-month-old WT female mice and placed in 0 mM Ca^2+^ Ringer’s solution. Excessive connective tissue, tendons, fat, blood clots and vessels were removed as much as possible. After cleanup, the muscles were transferred into the digestion media (DMEM containing 5% FBS, 1% Pen/Strep and 10 mM HEPES) and minced into small blocks using scissors. Collagenase II (Worthington LS004176, ≥125 units/mg), dispase II (Sigma D46931G, ≥0.5 units/mg), hyaluronidase (Worthington LS002592, ≥300 USP/NF units/mg) and Dnase I (Worthington LS002139, ≥2,000 Kunitz units/mg) were added at the final concentration of 0.26%, 0.24%, 0.16% and 0.04% (weight/volume), respectively. The digestion system was placed in an orbital shaker running at 70 rpm at 37 °C for 45 min. The digested muscles were triturated 10 times with a 20-gauge needle attached to a 5 ml syringe. Afterwards the triturated mixture was pipetted onto a pre-wetted 100 μm strainer and centrifuged at 100 x g for 2.5 min to precipitate the bulky myofiber debris. The supernatant was transferred onto a pre-wetted 40 μm strainer, and cells were collected by centrifuged at 1200 x g for 6.5 min. After the removal of supernatant, cells were resuspended with 5 ml growth medium (Ham’s F-10 medium + 20% FBS + 1%Pen/Strep + 5ng/ml bFGF + 10 μg/ml Plasmocin and pre-plated in non-coated T25 flasks at 37 °C for 30 min. The unattached cells were transferred together with medium to Matrigel coated T25 flasks and cultured till reaching about 50% confluence. Pre-plating continued for three more passages to diminish non-myoblast cells.

### Chemical and AAV treatment of cultured myotubes

Cultured myoblasts derived from soleus or EDL muscles of 4-month-old WT mice (seeded in laminin coated chambered cover glass at about 2.5x10^4/compartment) were induced to differentiate into myotubes in low serum medium (DMEM+2% horse serum) for four days, with medium changed every other day. 4-HNE treatment was performed at 0, 7.5, 15, 30 μM for 2 hours. The treatment time was determined based on a previous study showing 4-HNE notably induced oxidative stress as early as 2 hours in Swiss 3T3 fibroblasts [22].

AAVMYO(9P1)-tMCKp-huAldh3a1.HA-WPRE (AAV-Aldh3a1 for short) was produced by Vector Biolabs. AAVMYO(9P1) is a mutant of AAV9 that shows superior efficiency and specificity in skeletal and cardiac muscles [86]. The triple muscle creatine kinase promoter (tMCKp) specifically drives gene expression in differentiated muscle tissue [87]. To carry out AAV transduction, cultured myoblasts derived from soleus or EDL muscles of WT mice 4-5 months of age (seeded in laminin coated 48-well plate or chambered cover glass at about 2.5x10^4/compartment) were induced to differentiate into myotubes with low serum medium containing AAV-Aldh3a1 (1x10^11 GC/ml) for 2 days and regular differentiation medium for another 2 days. 4-HNE treatment (30 μM for 2 hours) was performed at the end of the second day. After the transient exposure to 4-HNE, myotubes were cultured for another 16 hours in regular low serum medium before TUNEL assay (Biotium 30074) or LDH leakage-based cytotoxicity assay (Promega G1780). This is because apoptosis can take 12-24 hours to occur [48].

### TUNEL assay

Triplicate cultures of myotubes in chambered cover glass (Cellvis C8-1.5H-N) were fixed and permeabilized with precooled methanol at -20 °C for 15 min. Afterwards, the samples were rehydrated by washing with PBS for 3 times (5 min each). TUNEL staining was performed following the manual of Biotum CF640R TUNEL Assay Apoptosis Detection Kit. The incubation time with the reaction mix (containing TdT enzyme) was 1.5 hours at 37 °C. Afterwards, the samples were washed twice with PBS and incubated with blocking buffer for 45 min at room temperature, followed by immunostaining with Aldh3a1 and myosin heavy chain (DSHB, MF-20 1:100) antibodies. Counterstaining with DAPI was done to calculate the ratio of nuclei with fragmented genomic DNA.

### Non-radioactive cytotoxicity (LDH leakage) assay

We generally followed the protocol of the assay kit (Promega, G1780). In brief, 5-6 replicates of myotube cultures with or without AAV transduction, 4-HNE treatment, were cultured in 300 μl regular low serum medium for 16 hours, along with wells containing only 300 μl medium but no cells (to serve as medium only controls and volume correction controls). Next morning, 50 μl of culture supernatant was collected and 50 μl of lysis solution (contains Triton X-100) was added to lyse all the remaining myotubes (37 °C, 45 min). The supernatant, the medium only control, the all-cell-lysed solution (12.5 μl diluted to 50 μl with PBS+1% BSA, 1:4 dilution) and the volume correction controls (medium only wells with 50 μl replaced with lysis solution and diluted 1:4 with PBS+1% BSA) were combined with 50 μl of chromogenic substrates of lactate dehydrogenase (LDH, a stable cytosolic enzyme that is released upon cell death), respectively, for absorbance measurement by a plate reader (SpectraMax i3x, Molecular Devices). Absorbance at 490 nm was recorded every 5 min for 35 min for each sample. The largest rate of absorbance increase (Vmax) calculated from 3 continuous points out of the 7 time points measured reflects the concentration of LDH. To compare the percentages of dead myotubes under different treatment conditions, we calculated the percentage of LDH in the culture supernatant to total LDH as follows: 100*(Vmax(supernatant)-Vmax(medium only control))/(4*(Vmax(all-cell-lysed)-Vmax(volume correction control))+(Vmax(supernatant)-Vmax(medium only control))/6).

### Transfection of GFP-MG53 and recording of saponin induced MG53 membrane translocation

4.5x10^4 of myoblasts derived from soleus muscles of 4-month-old WT mice were seeded into glass bottom dishes (Matsunami D35-14-1.5-U) and cultured in growth medium overnight before transfection of 3 μg GFP-MG53 plasmids with TransfeX transfection reagent (ATCC ACS-4005). We selected soleus derived myoblasts because they form multi-nuclei, lengthy myotubes more robustly than EDL derived ones, facilitating the measurement of membranous GFP-MG53 signals.GFP-MG53 was a gift from Dr. Jianjie Ma’s lab [34]. One day after transfection, the culture medium was changed to low serum medium to induce differentiation for 4 days, with medium changed every two days. AAV- Aldh3a1 treatment group was incubated with the AAV vector at the first two days of differentiation. 4- HNE treatment group was incubated with 30 μM 4-HNE for 2 hours at the end of the 4-day differentiation. The myotubes were washed three times with 0 mM Ca^2+^ Ringer’s solution containing 0.5 mM EGTA (Sigma Aldrich E4378) and 90 μM BTS (Sigma Aldrich 203895) to prevent contraction.

Time lapse imaging of myotubes were performed with Leica TCS SP8 confocal microscope at an interval of 2 sec for 3 min. 0.00375% saponin (Thermo Scientific A18820.14) was applied 10 sec after recording started. Regions of interest were generated at plasma membrane regions and background signals from myotube free area was deduced to acquire GFP-MG53 intensity profile (*F*) over time. *F_0_* was the averaged intensity of the first 4 frames.

### Data analysis and statistics

Box-and-dot plots were generated with ggplot2 package of R (version 4.2.3). The lower hinge, median line and upper hinge correspond to the first, second and third quartiles. The lower and upper whiskers extend from the hinges to the largest value no further than 1.5 times of inter-quartile range (distance between the first and third quartiles). Data beyond the end of the whiskers are outlying points. The stat_compare_means function of ggplot 2 was used to compare the means of two groups. Wilcoxon rank-sum test was used unless there were tie values between two groups (**Figure 8C**) or the group had less than 4 samples (**Figure 9A**). In those cases, t-test was used instead. For multi- group data, one-way ANOVA P values were generated by aov function of R.

## Acknowledgement

We thank Dr. Yongfu Wang at and Dr. Ji Pang, who previously worked at Stowers Institute for Medical Research, for the help in trouble-shooting the glyoxal fixation protocol for section immunofluorescence. This study has been supported by grants from NIH (R01NS105621 to JZ, R01NS129219 to JZ and JM) and the Department of Defense AL170061(W81XWH1810684) to JZ.

## Author contributions

Ang Li, Conceptualization, Data curation, Formal analysis, Validation, Investigation, Visualization, Methodology, Writing – original draft, Project administration, Writing – review and editing; Li Dong, Xuejun Li, Jianxun Yi, Data curation, Validation, Investigation, Methodology; Jianjie Ma, Conceptualization, Funding acquisition, Writing – review and editing; Jingsong Zhou, Conceptualization, Resources, Supervision, Funding acquisition, Validation, Investigation, Visualization, Methodology, Writing – original draft, Project administration, Writing – review and editing

## Abbreviations

4-HNE: 4-hydroxy-2-nonenal
AChR: acetylcholine receptor
ALS: amyotrophic lateral sclerosis
EDL: extensor digitorum longus
SOL: soleus
DIA: diaphragm
EOM: extraocular muscle
ELISA: enzyme-linked immunosorbent assay
HL: hindlimb
MDA: malondialdehyde
NMJ: neuromuscular junction
SC: satellite cell
SNT: sciatic nerve transection
SFN: sulforaphane

---

**Video 1.** Time lapse imaging of GFP-MG53 transfected myotubes without 4-HNE or AAV-Aldh3a1 treatment.

**Video 2.** Time lapse imaging of GFP-MG53 transfected myotubes treated with 30 μM 4-HNE for 2 hours before recording.

**Video 3.** Time lapse imaging of GFP-MG53 transfected, AAV-Aldh3a1 transduced myotubes treated with 30 μM 4-HNE for 2 hours before recording.

**Figure 1-figure supplement 1.**
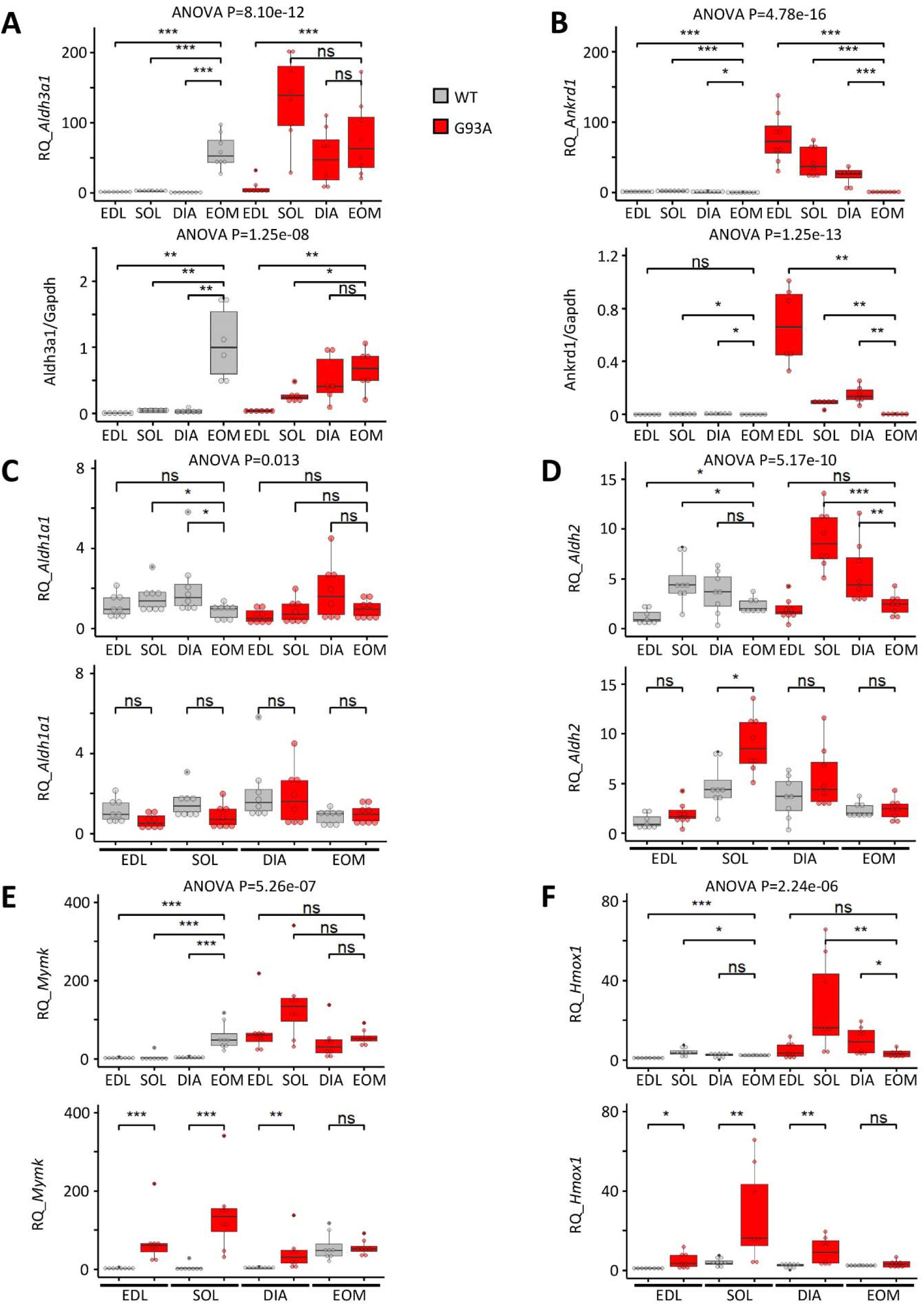
Additional qRT-PCR and Western blot quantification results for different muscles from end-stage G93A mice and WT littermates. (**A**) qRT-PCR (upper) and Western blot (lower) relative quantification results of *Aldh3a1* (against housekeeping gene *Gapdh*) in EDL, soleus (SOL), diaphragm (DIA) and extraocular muscles (EOM) from end-stage G93A mice and WT littermates. (**B**) qRT-PCR (upper) and Western blot (lower) relative quantification results of *Ankrd1* in different muscles from end-stage G93A mice and WT littermates. (**C-F**) qRT-PCR relative quantification results of *Aldh1a1*, *Aldh2*, *Mymk* and *Hmox1*. *** P < 0.001; ** P < 0.01; * P < 0.05; ns, not significant (Wilcoxon rank-sum test). ANOVA P values are also shown. N = 8 (4 pairs of male mice and 4 pairs of female mice). RQ, relative quantification. Please also see **Figure 1-Source Data 1**.

**Figure 1-figure supplement 2.**
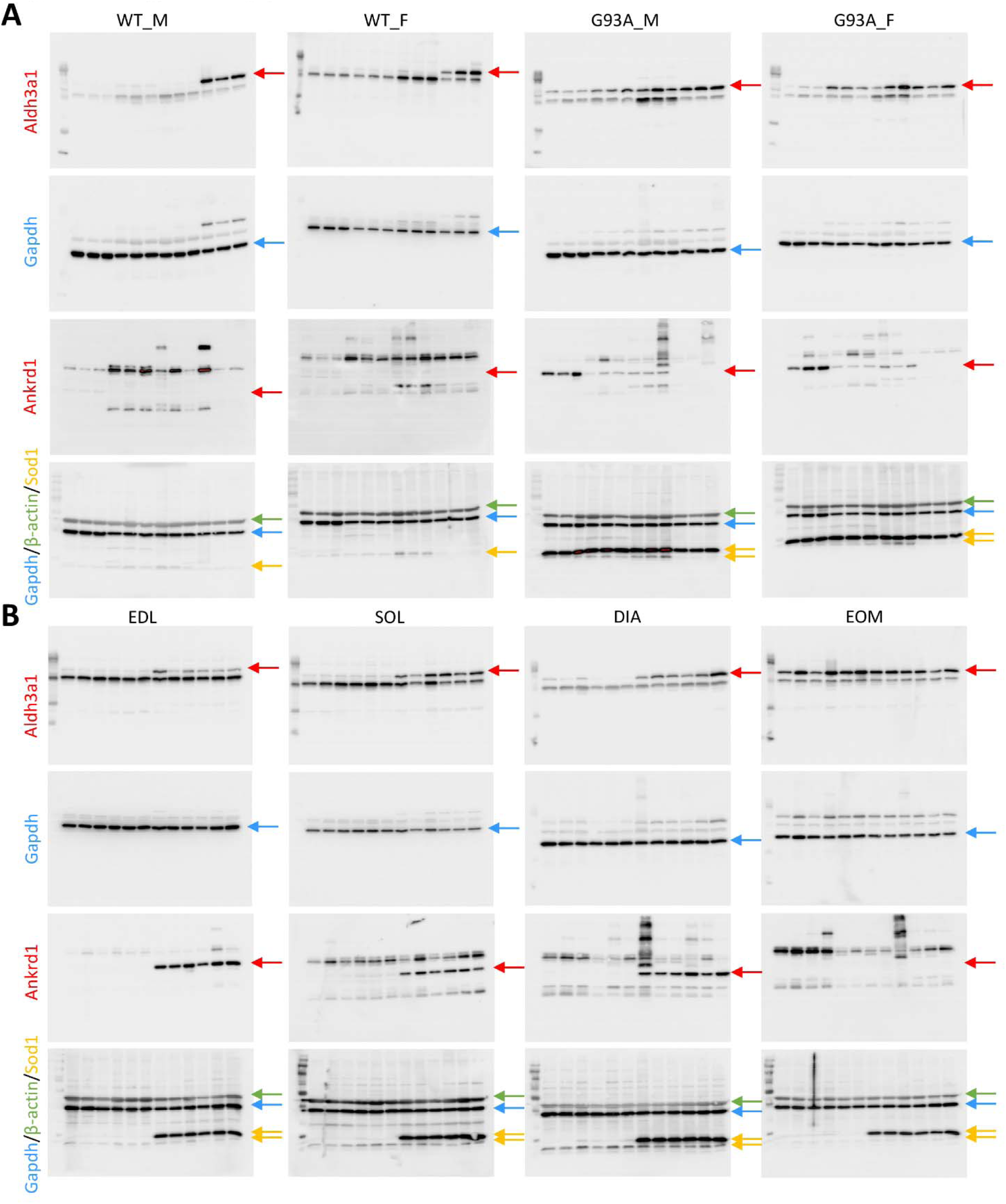
Original images for Western blots in. Figure 1. (**A**) Original images for Western blots in Figure 1B. Red arrows indicate Aldh3a1 (51 kDa) or Ankrd1 (36 kDa) bands. Blue arrows indicate Gapdh (36 kDa) bands. Green arrows indicate β-actin (42 kDa) bands. Orange arrows indicate hSod1 (16-20 kDa) bands (the upper band is human Sod1^G93A^, the lower band is mouse endogenous Sod1). (**B**) Original images for Western blots in Figure 1C.

**Figure 3-figure supplement 1.**
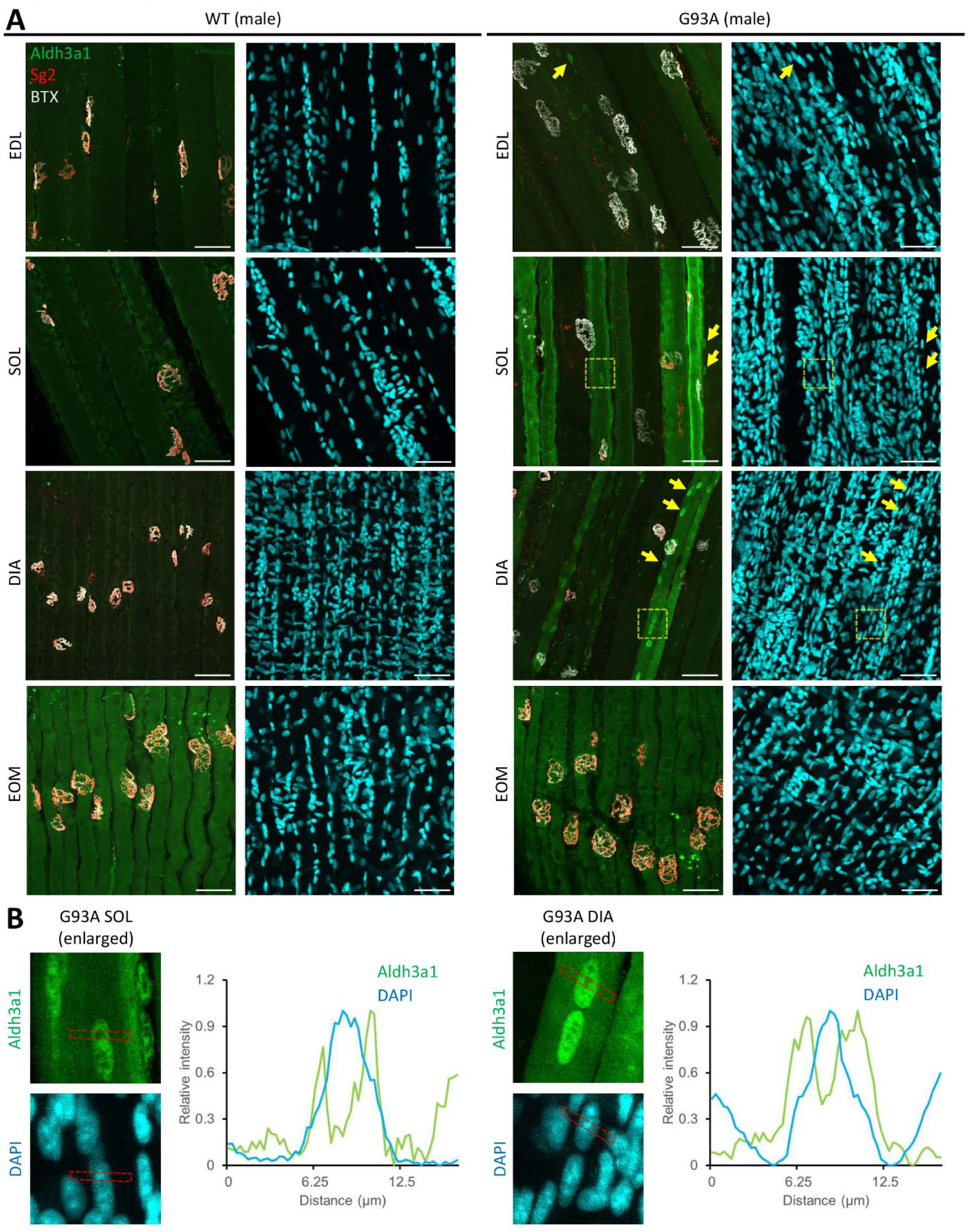
Additional images of Aldh3a1 whole-mount immunostaining in different muscles from end-stage G93A mice and WT littermates. (**A**) Representative compacted z-stack scan images of whole-mount EDL, soleus diaphragm extraocular muscles stained with antibodies against Aldh3a1, Sg2 (labeling axon terminals), Alexa Fluor conjugated α-Bungarotoxin (BTX, labeling AChRs on muscle membrane) and DAPI (labeling nuclei). N = 6 (3 pairs of male and 3 pairs of female). Scale bars, 50 μm. Yellow arrows highlight nuclei with Aldh3a1 enrichment. Dashed yellow boxes denote regions enlarged in Panel B for kymographic measurement. (**B**) Profiling the relative intensity of Aldh3a1 and DAPI fluorescent signals along the strips denoted by the red boxes. Relative intensities are calculated as (*F* − *F_min_*)/(*F_max_* − *F_min_*).

**Figure 4-figure supplement 1.**
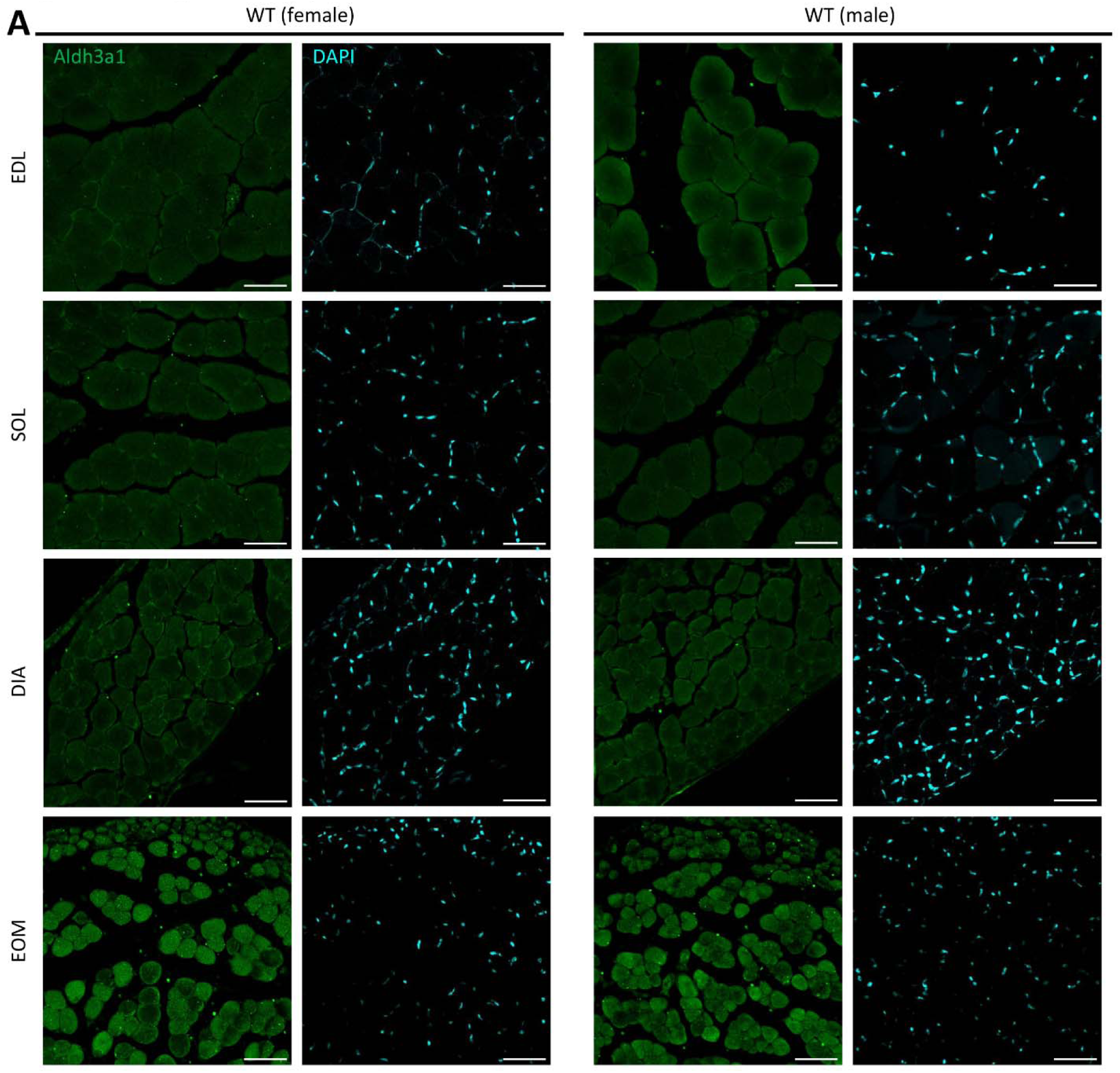
Section immunostaining results of Aldh3a1 in different muscles from WT mice 4-5 months of age. (**A**) Transverse sections of EDL, soleus, diaphragm and EOMs from WT mice stained with antibodies recognizing Aldh3a1. N = 6 (3 male and 3 female). Scale bars: 50 μm.

**Figure 5-figure supplement 1.**
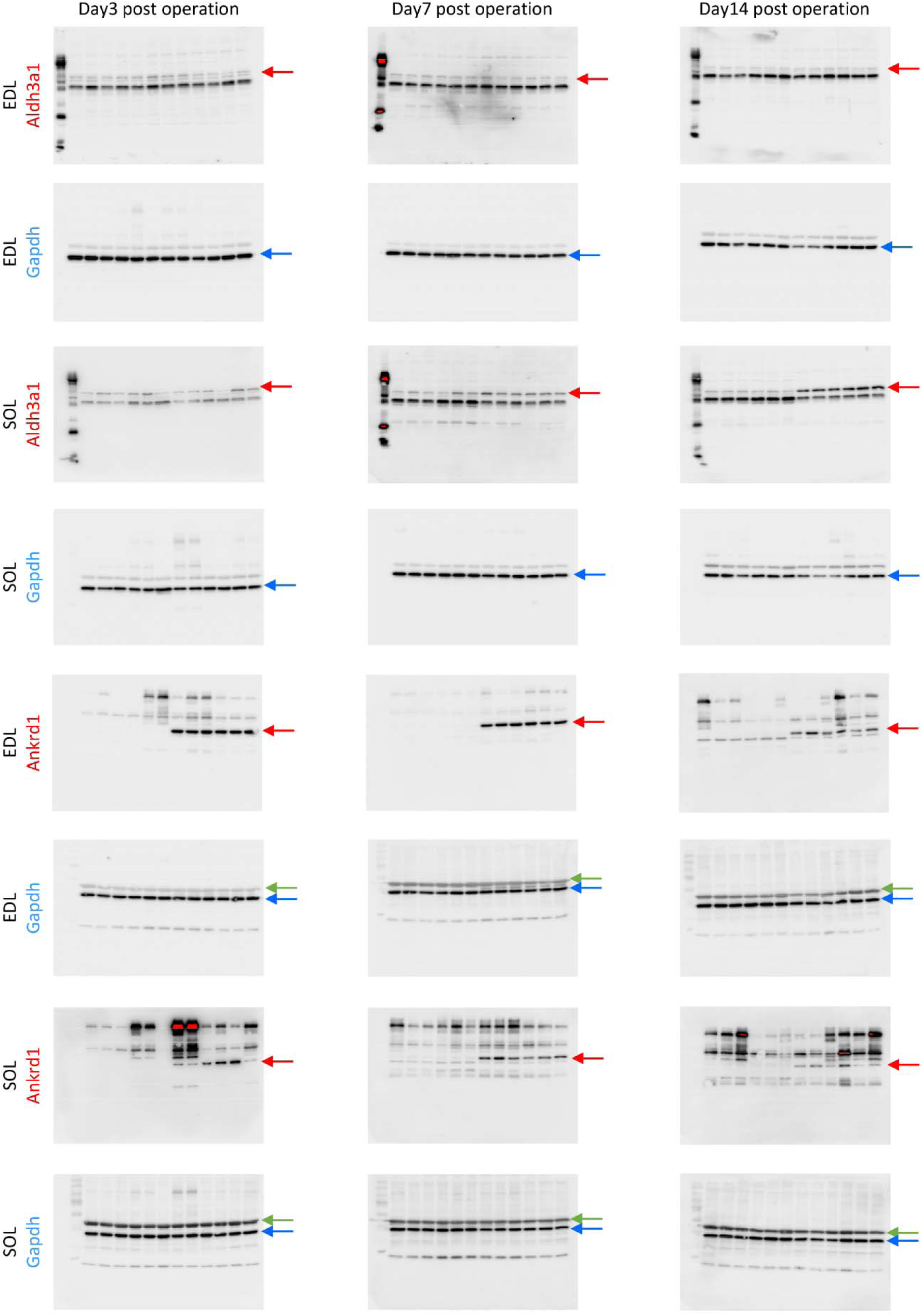
Original images of Western blots in Figure 5. Red arrows indicate Aldh3a1 (51 KDa) or Ankrd1 (36 kDa) bands. Blue arrows indicate Gapdh (36 kDa) bands. Green arrows indicate β-actin (42 kDa) bands.

**Figure 7-figure supplement 1.**
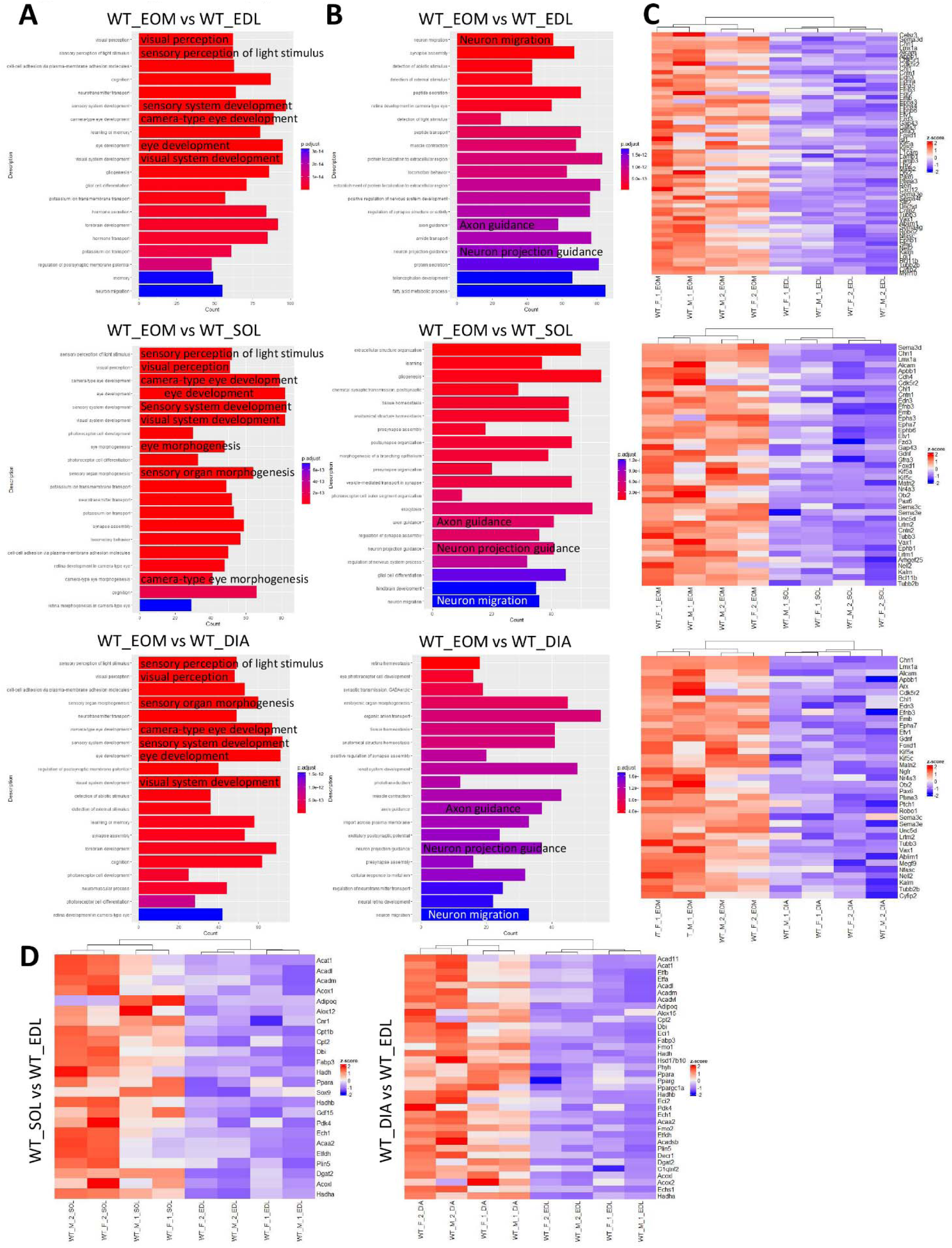
Gene ontology analysis and heatmaps of genes differentially expressed between different groups of muscles in WT mice. (**A**) Top 20 biological processes in gene ontology analysis for genes expressed higher in EOMs than other muscles from WT mice. Processes related to eye development are denoted. (**B**) Axon guidance/neuron migration related biological processes highlighted by gene ontology analysis for genes expressed higher in EOMs than other muscles from WT mice. (**C**) Heatmaps of axon guidance related DEGs comparing EOMs to other muscles from WT mice. (**D**) Heatmaps of lipid oxidation related DEGs comparing soleus and diaphragm to EDL muscles from WT mice.

**Figure 1-Source Data 1. qRT-PCR results for *Aldh1a1, Aldh2, Aldh3a1, Ankrd1, Mymk* and *Hmox1* expression levels in whole muscles of different origins.**

**Figure 1-Source Data 2. Western blot quantification results for Aldh3a1 and Ankrd1 protein levels in whole muscles of different origins.**

**Figure 5-Source Data 1. qRT-PCR results of the fold of change of *Aldh3a1* and *Ankrd1* expression levels in whole muscles with SNT compared to the sham operated controls.**

**Figure 5-Source Data 2. Western blot quantification results of the fold of change of Aldh3a1 protein levels in whole muscles with SNT compared to the sham operated controls.**

**Figure 5-Source Data 3. Western blot quantification results of the fold of change of Ankrd1 protein levels in whole muscles with SNT compared to the sham operated controls.**

**Figure 7-Source Data 1. Entrez IDs, symbols, log2 fold changes and adjusted p values of the 866 EOM signature genes.**

**Figure 7-Source Data 2. qRT-PCR results for *Hmox1* and *Aldh3a1* expression levels in EDL and soleus-SC derived myotubes with and without SFN treatment.**

**Figure 8-Source Data 1. 4-HNE levels in EDL and soleus muscles from end-stage G93A mice and age-matched WT controls measured by ELISA. .**

**Figure 8-Source Data 2. Quantification results of the percentages of TUNEL positive nuclei in myotubes treated with different concentrations of 4-HNE.**

**Figure 8-Source Data 3. Quantification results of LDH leakage percentages of myotubes treated with different concentrations of 4-HNE.**

**Figure 9-Source Data 1. Western blot quantification results for Aldh3a1 protein levels in myotubes with or without transduction of AAV-Aldh3a1.**

**Figure 9-Source Data 2. Quantification results of the percentages of TUNEL positive nuclei in myotubes with or without transduction of AAV-Aldh3a1 before 4-HNE treatment.**

**Figure 9-Source Data 3. Quantification results of LDH leakage percentages of myotubes with or without transduction of AAV-Aldh3a1 before 4-HNE treatment.**

**Figure 10-Source Data 1. Ratios between maximum *F* and *F_0_* in each recorded myotubes transfected with GFP-MG53 and whether the *F_180_* is higher or lower than *F_0_*.**

